# Dietary protein governs the role of insulin signaling in the postprandial regulation of hepatic mTORC1

**DOI:** 10.1101/2025.10.01.679861

**Authors:** Krystle C. Kalafut, Yann Cormerais, Madi Y. Cissé, Samuel C. Lapp, Karen E. Inouye, Gökhan S. Hotamışlıgil, Brendan D. Manning

**Affiliations:** Department of Molecular Metabolism, Harvard T.H. Chan School of Public Health, Boston, MA, USA; Department of Cell Biology, Harvard Medical School, Boston, MA, USA

## Abstract

The nutrient-sensing mechanistic target of rapamycin complex 1 (mTORC1) signaling pathway controls cellular and organismal growth and metabolism, while aberrant activation is linked to human disease, including metabolic disease. Cellular studies have established several regulatory mechanisms influencing mTORC1 activation, but the physiological signals that control mTORC1 at the organismal and tissue levels are less well defined. mTORC1 is dynamically regulated by fasting and feeding in metabolic tissues, with both nutrients and insulin proposed to activate mTORC1 in response to feeding. Here, a liver-specific mouse model that disconnects mTORC1 activation from AKT-mediated TSC2 phosphorylation is employed. This genetic mouse model demonstrates that AKT-mediated TSC2 phosphorylation is the predominant mechanism of hepatic mTORC1 induction by insulin but is dispensable for activation by feeding. Furthermore, dietary protein is critical and dictates the insulin-responsiveness of hepatic mTORC1 signaling. Contrary to dogma, hepatic mTORC1 signaling was not elevated in response to diet-induced obesity associated with the phenotypes of type-2 diabetes, including hyperinsulinemia, systemic insulin resistance, and hyperglycemia, and blocking hepatic AKT-TSC-mTORC1 signaling did not prevent these metabolic impairments. Evidence is also provided supporting a role for glucagon in hepatic mTORC1 suppression during fasting. This study reveals a hierarchy of physiological signals regulating hepatic mTORC1.

## Introduction

The liver is a central hub for macronutrient metabolism that coordinates physiological responses to dynamic nutrient and energy states, such as across physiological fasting and feeding cycles^1, 2^. To coordinate these metabolic responses, the liver must be able to both monitor and adapt to acute changes in nutrient or energy availability. A key function of the liver is the regulation of glucose and lipid homeostasis in response to hormonal cues, such as that from insulin^3^. Importantly, hepatic insulin resistance is implicated in metabolic disease, including obesity and type 2 diabetes^4^. However, the relevant targets of insulin signaling in the liver and the associated contribution to hepatic or systemic metabolism remain incompletely understood.

At the cellular level, mechanistic target of rapamycin complex 1 (mTORC1) signaling controls the balance between anabolic and catabolic metabolism^5^. mTORC1 activation promotes anabolic processes and biosynthetic pathways in response to a variety of nutrients and signals propagated by growth factors and hormones, tailoring growth and metabolism to local and systemic nutrient status^5^. In vitro studies have established insulin, as well as glucose and amino acids, as a robust stimulus of mTORC1 activation^6^. In the liver, mTORC1 activation is sensitive to fluctuations in nutrients, with acute activation upon feeding and suppression during fasting^7, 8^, and aberrant hepatic mTORC1 activation has been linked to metabolic disease^9–11^. However, it is unclear how specific upstream inputs cooperate to regulate mTORC1 in physiological or pathological settings. Therefore, dissecting the mechanisms of hepatic mTORC1 regulation and its consequences for systemic metabolism is central to understanding metabolic control in health and disease.

Insulin activates mTORC1 via PI3K-AKT signaling, and several mechanisms of AKT-dependent mTORC1 activation have been identified in cell culture models^12–17^. The most well-characterized mechanism involves AKT-mediated inhibitory phosphorylation of the TSC2 component of the TSC complex, a key regulatory hub that inhibits mTORC1^18–20^. TSC2 is a GTPase activating protein (GAP) for the small GTPase Rheb, which facilitates maximal activation of the mTOR protein kinase by engaging mTORC1 at the lysosomal surface specifically when GTP-bound^21–25^. Therefore, the TSC complex inhibits mTORC1 activation by promoting the conversion of GTP-to GDP-bound Rheb. In response to insulin, AKT phosphorylates TSC2 at five distinct residues corresponding to Ser939, Ser981, Ser1130, Ser1132, and Thr1462 of the full-length human TSC2 protein, and this regulation is required for subsequent mTORC1 activation in cell culture models^18–20, 24, 26^. AKT-mediated phosphorylation of TSC2 inhibits the TSC complex by causing its dissociation from the lysosome, where a subpopulation of Rheb resides, thereby allowing Rheb-GTP to locally accumulate and activate mTORC1^24^. The five AKT-mediated phosphorylation sites are conserved among vertebrate TSC2 orthologs, suggesting their physiological significance in complex multicellular organisms, but this has not been directly tested in mice^27^.

The TSC complex is essential for the dynamic regulation of mTORC1 in the liver, as mice with liver-specific ablation of TSC complex components fail to suppress mTORC1 activation during fasting^7, 8, 28, 29^. However, multiple signaling pathways converge on the TSC complex to either promote or suppress mTORC1 activity^27^, thus it is not entirely clear which signal is disrupted in such models. Like genetic loss of TSC complex components, other liver-specific mouse models used to study mTORC1 regulation *in vivo*, including genetic ablation of AKT or of mTORC1 components^7, 8, 28, 30–37^, disrupt the entire, highly branched signaling network and may lead to compensatory rewiring not reflective of physiological signaling^5, 6^. Thus, more precise models are needed to understand the physiological regulation of mTORC1 and its implications for health and disease.

In addition to regulation by growth factors through the TSC complex and Rheb, mTORC1 activation is also sensitive to cellular amino acid levels through a pathway regulating a separate set of small G-proteins, the Rag GTPases, that control mTORC1 lysosomal localization^38^. In nutrient-replete conditions, Rag heterodimers consisting of RagA-or RagB-GTP (RagA/B-GTP) complexed with RagC- or RagD-GDP (RagC/D-GDP) recruit mTORC1 to the lysosome, where it can access Rheb-GTP^38, 39^. Insight from cell culture studies supports a model whereby maximal mTORC1 activation requires the presence of both extracellular growth signals and sufficient intracellular nutrients to convert the Rheb and Rag GTPases, respectively, to the correct guanylate binding state to engage mTORC1 at the lysosomal surface^5^. Importantly, like liver-specific loss of the TSC complex, mice that expresses a GTP-bound mutant of RagA exhibit aberrant hepatic mTORC1 signaling even during fasting^40^, demonstrating that signals through both the TSC-Rheb and Rag pathways are critical for the proper dynamic regulation of mTORC1 with fasting and feeding.

Given its role in metabolic regulation, hepatic mTORC1 signaling has been proposed to contribute to impaired glucose and lipid homeostasis in obesity and diabetes^6, 9, 10, 41^. Some studies have concluded that hepatic mTORC1 is chronically elevated in obese mice, and mouse models characterized by repression of hepatic mTORC1 signaling exhibit protection from obesity-associated metabolic impairments ^9–11, 37, 42^. However, the mechanisms driving hepatic mTORC1 activation in obesity and the causal link of hepatic mTORC1 to metabolic dysfunction have not been clearly defined. It has been widely proposed that chronically active mTORC1 can contribute to insulin resistance through negative feedback regulation of insulin signaling to AKT^10, 43–46^. This mechanism has been supported by in vivo studies showing that liver AKT activation by feeding or insulin is severely blunted in mice with chronic activation of hepatic mTORC1 and enhanced in mice lacking hepatic mTORC1 or its direct substrate S6K1^7, 9, 11, 28, 47^. Importantly, enhanced liver AKT activation in obese *S6K*^-/-9^ or liver-specific S6K knockdown mice^11^ is accompanied by improved systemic insulin tolerance and insulin-mediated suppression of hepatic glucose production^11^. Thus, defining the mechanisms underlying hepatic mTORC1 regulation by pathological signals, such as hyperinsulinemia and overnutrition, will be necessary to understand its putative role in metabolic diseases.

Here, we describe a genetic mouse model that disconnects hepatic insulin signaling from mTORC1 activation. We utilize mice that conditionally expresses wild-type human TSC2 (TSC2-WT) or a TSC2 phospho-mutant that cannot be phosphorylated by AKT (TSC2-5A) in place of endogenous TSC2. Hepatic mTORC1 signaling is unresponsive to insulin but activated normally by feeding in liver-specific TSC2-5A (L-TSC2-5A). We find that dietary protein is the dominant factor driving hepatic mTORC1 activation by feeding and that dietary protein content influences the role of insulin-AKT-TSC signaling in mTORC1 activation—a hierarchy that may be unique to the liver. In contrast to previous literature^9, 10^, high-fat diet-induced obesity, hyperinsulinemia, and diabetes were not accompanied by elevated mTORC1 signaling in liver tissue. Furthermore, L-TSC2-5A mice were not protected from obesity-associated impairments in glucose homeostasis, despite mTORC1 signaling remaining unresponsive to insulin in this model. We also investigate glucagon as a negative regulator of hepatic mTORC1 and provide genetic evidence that glucagon signaling contributes to mTORC1 suppression during fasting. These data provide key insights into physiological regulators of mTORC1 and highlight the importance of understanding distinct mechanisms of nutrient-sensing across metabolic tissues.

## Results

### Genetic partitioning of AKT-TSC signaling from hepatic mTORC1 activation does not impair basal mTORC1 activation or systemic metabolic health

Previous reports from our lab and others have demonstrated that the TSC complex is required for the dynamic regulation of hepatic mTORC1 in response to fasting and feeding^7, 8, 28^. As expected, 1 h-refeeding after an overnight fast induced phosphorylation of AKT, as well as the mTORC1 targets S6K1 and 4EBP1 (indicated by mobility shift), in the mouse liver (**Figure 1A**). Importantly, feeding also induced phosphorylation of TSC2 at T1462, one of the established sites regulated by AKT (**Figure 1A**)^18, 20^, indicating that AKT-mediated TSC2 phosphorylation coincides with mTORC1 activation in physiological settings.

**Figure 1.**
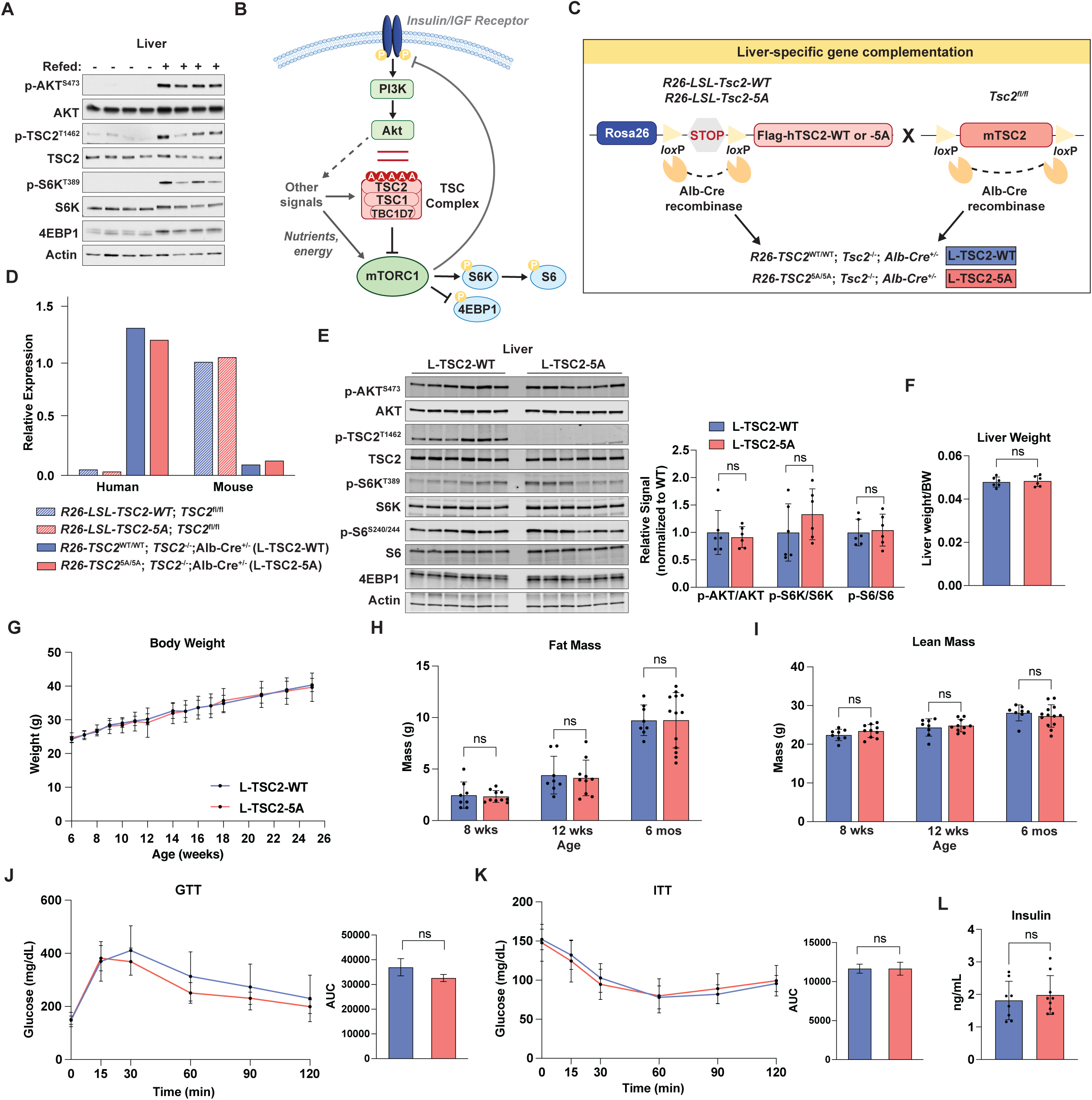
Genetic partitioning of AKT-TSC signaling from hepatic mTORC1 activation does not impair basal mTORC1 activation or systemic metabolic health. (A) Eight-week-old male C57BL/6J mice were fasted overnight for 16 h and, where indicated, refed for 1 h prior to immunoblot analyses of whole liver extracts; n = 4/group. (B) Schematic depicting activation of mTORC1 by insulin-AKT signaling, highlighting the TSC2-5A phosphomutant characterized in this study. (C) Schematic of the conditional L-TSC2-WT and -5A genetic mouse models. The human TSC2-WT or -5A transgene expressed from the *Rosa26* locus replaces endogenous mouse *Tsc2* expression following Cre-mediated recombination. Liver-specific TSC2-WT or -5A expression was achieved through Cre-driven expression from the Albumin (Alb) promoter. (D) Recombination at the *Rosa26* and *Tsc2^fl/fl^* gene loci was validated by RT-qPCR analysis of human versus mouse *Tsc2* expression in liver tissue in mice with or without expression of Albumin-Cre. (E) Immunoblots of liver lysates from 12-week-old L-TSC2-WT or -5A male mice fed ad libitum with accompanying bar graphs of the mean ± SD of phospho-to-total protein signal quantifications normalized to the L-TSC2-WT group; n = 6/group. (F) Liver weights (normalized to body weight) of 12-week-old male L-TSC2-WT and -5A mice fed ad libitum graphed as mean ± SD (n = 6 /group). (G) Body weights of male L-TSC2-WT and -5A mice up to 6 months of age graphed as mean ± SD; n=8-13/group/timepoint. There were no significant differences in body weight between genotypes. (H, I) Fat (H) and lean (I) mass of L-TSC2-WT (blue) and -5A (red) male mice measured by EchoMRI at indicated ages graphed as mean ± SD; n = 8-13/group. (J) Glucose tolerance test performed on 6-month-old L-TSC2-WT (blue) and -5A (red) mice after a 6-h daytime fast followed by i.p. injection of glucose (2g/kg). Blood glucose measurements over time are plotted as mean ± SD. Area under the curve (AUC) values are graphed as mean ± SEM; n = 8 WT, 9 5A. (K) Insulin tolerance test performed on 6-month-old L-TSC2-WT (blue) and -5A (red) mice after a 6-h daytime fast followed by i.p. injection of insulin (0.5 U/kg). Blood glucose measurements over time are plotted as mean ± SD. AUC values are graphed as mean ± SEM; n=7 WT, 10 5A. (L) Plasma insulin following 6-h daytime fast in 6-month-old male L-TSC2-WT (blue) and -5A (red) mice graphed as mean ± SD; n = 8 WT, 9 5A. Statistical analysis for: (E, F, and L) Welch’s t-test; (G) mixed-effects analysis with Šidák correction for multiple comparisons; (H and I) two-way ANOVA with Šidák correction; (J and K) two-way ANOVA with Šidák correction for curves, Welch’s t-test for AUC. Not significant (ns = p≥0.05)

To determine the causal role of AKT-mediated TSC2 phosphorylation for the control of mTORC1 signaling in mammalian tissues, we have generated a knock-in mouse model expressing wild-type TSC2 (TSC2-WT) or TSC2 with alanine mutations at the five AKT-regulated phosphorylation sites (TSC2-5A)^48^ (**Figure 1B**). The transgenes are expressed conditionally from the *Rosa26* (R26) locus through the use of a transcriptional and translational stop cassette flanked by LoxP sites (LSL) and encode Flag-tagged TSC2-WT or -5A human cDNAs^48, 49^ (**Figure 1C**). Thus, expression from the ubiquitous R26 promoter is controlled by Cre recombinase, allowing transgene expression with concomitant deletion of endogenous *Tsc2* when bred to *Tsc2^fl/fl^* mice (**Figure 1C**). *R26-LSL-Tsc2^WT/WT^; Tsc2^fl/fl^*and *R26-LSL-Tsc2^5A/5A^; Tsc2^fl/f^*^l^ mice were crossed to mice expressing Cre recombinase from the hepatocyte-specific Albumin promoter (*Alb-Cre^+/-^*) to generate mice with liver-specific expression of the TSC2-WT or -5A transgenes coupled to liver-specific deletion of endogenous *Tsc2* (referred to throughout as L-TSC2-WT or L-TSC2-5A) (**Figure 1C**). Cre-dependent recombination at both the *R26-LSL* and endogenous *Tsc2* loci in liver tissue was confirmed by RT-qPCR analysis of mouse versus human *Tsc2* expression (**Figure 1D**).

An analysis of signaling in liver tissue from L-TSC2-WT and -5A mice under ad libitum-fed conditions confirmed complete loss of phosphorylation of TSC2 by Akt at T1462 in the L-TSC2-5A livers. However, there were no significant differences in the activating phosphorylation of AKT or in markers of mTORC1 signaling, including phosphorylation of S6K1, S6, and 4EBP1 between L-TSC2-WT and -5A livers (**Figure 1E**). Liver tissue weight, which is changed in mice with liver-specific disruption of the TSC complex or mTORC1^8^, was similar between L-TSC2-WT and -5A mice (**Figure 1F** (males); **Figure S1A** (females)). Unlike the whole-body TSC2-WT and -5A mice^48^, there was no difference between the liver-specific TSC2-WT and -5A mice in body weight and body composition measured at regular intervals from 6 weeks to 6 months of age, for males (**Figures 2G-I**) or females (**Figures S1B-D**). Furthermore, there was no difference in glucose or insulin tolerance or plasma insulin levels between age-matched L-TSC2-WT and L-TSC2-5A mice in either males (**Figures 1J-L**) or females (**Figures S1E-G**) at 6 months of age. These results indicate that AKT-mediated TSC2 phosphorylation is not required for basal mTORC1 activation in the liver or for the maintenance of glucose homeostasis on a normal chow diet.

**Figure 2.**
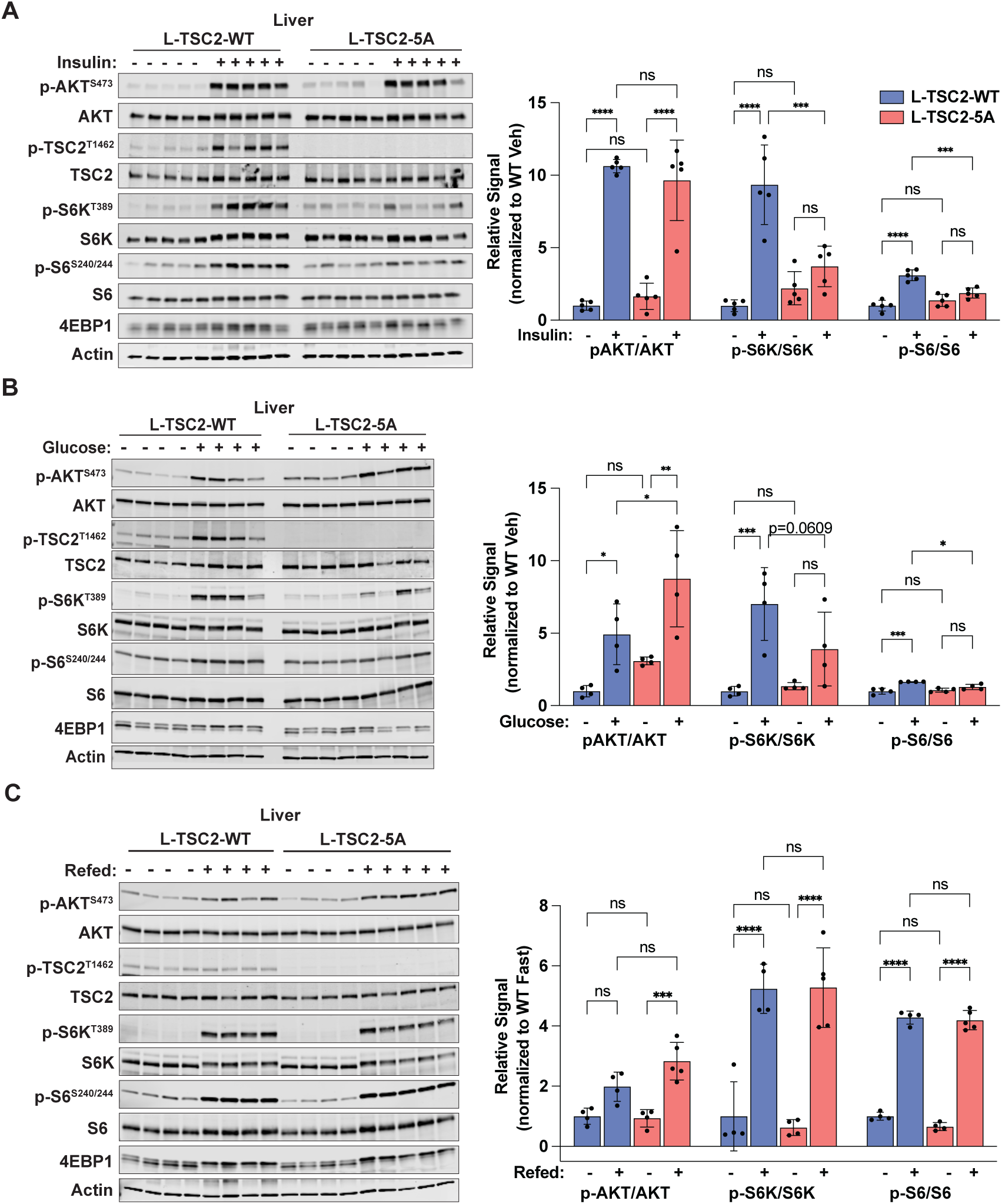
Hepatic mTORC1 activation by insulin or glucose, but not feeding, requires AKT-mediated TSC2 phosphorylation. (A) Immunoblots of liver lysates from 11-week-old L-TSC2-WT or -5A male mice fasted for 14 h overnight followed by i.p. injection with vehicle (saline) or insulin (0.5 U/kg) for 20 min with accompanying bar graphs of the mean ± SD of phospho-to-total protein signal quantifications normalized to the L-TSC2-WT vehicle-treated group; n = 5/group. (B) Immunoblots of liver lysates from 11-week-old L-TSC2-WT or -5A male mice fasted for 14 h overnight prior to administration of vehicle (saline) or glucose (3g/kg) by oral gavage for 25 min with accompanying bar graphs of the mean ± SD of phospho-to-total protein signal quantifications normalized to the L-TSC2-WT vehicle-treated group; n = 4/group. (C) Immunoblots of liver lysates from 10-week-old L-TSC2-WT or -5A male mice fasted for 14 h overnight and, where indicated, refed for one hour with accompanying bar graphs of the mean ± SD of phospho-to-total protein signal quantifications normalized to the L-TSC2-WT fasted group; n = 4-5/group. Statistical analysis for (A-C) two-way ANOVA with Šidák correction. Not significant (ns = p≥0.05), *p < 0.05, **p < 0.01, ***p < 0.001, and ****p < 0.0001.

### Hepatic mTORC1 activation by insulin or glucose, but not feeding, requires AKT-mediated TSC2 phosphorylation

AKT-mediated TSC2 phosphorylation is required for mTORC1 induction by insulin in cells^24^. To evaluate the role of this regulation in the intact liver, fasted L-TSC2-WT and L-TSC2-5A mice were treated with insulin or vehicle for 20 minutes, and AKT and mTORC1 signaling were assessed in liver tissue lysates. AKT was robustly activated by insulin in both L-TSC2-WT and -5A livers (**Figure 2A**). This was also reflected in phosphorylation of TSC2 at T1462 in L-TSC2-WT, but not L-TSC2-5A, livers. Importantly, mTORC1 stimulation by insulin was impaired in L-TSC2-5A compared to L-TSC2-WT livers (**Figure 2A**). As a control, signaling was also analyzed in gastrocnemius muscle from the same mice, where mTORC1 was stimulated by insulin to a similar extent in L-TSC2-WT and -5A mice (**Figure S2A**), confirming that the impaired mTORC1 induction in L-TSC2-5A mice is a liver-intrinsic effect. To determine whether L-TSC2-5A mice have deficient hepatic mTORC1 induction in response to endogenous insulin secretion, fasted L-TSC2-WT and L-TSC2-5A male mice were administered glucose or vehicle via oral gavage and signaling was assessed in liver tissue collected after 25 minutes. Oral glucose administration resulted in significant elevation in blood glucose and insulin compared to vehicle treatment, with similar levels in both L-TSC2-WT and -5A mice (**Figure S2B, C**). AKT was activated by glucose in both L-TSC2-WT and -5A livers, but the induction of mTORC1 signaling was blunted in L-TSC2-5A compared to L-TSC2-WT livers (**Figure 2B**). It is worth noting that, despite this decrease in mTORC1 signaling in the L-TSC2-5A liver, AKT activation was enhanced in response to glucose, perhaps reflecting partial relief of mTORC1-dependent feedback inhibition of insulin signaling in the L-TSC2-5A livers. These results confirm that TSC2 phosphorylation is necessary for proper hepatic mTORC1 induction by both exogenous and endogenous insulin and suggests that insulin signaling through AKT and TSC2 is the primary mechanism by which glucose acutely stimulates hepatic mTORC1.

The above experiments demonstrate that the L-TSC2-5A mouse model presents a genetic setting to evaluate the contribution of insulin signaling to hepatic mTORC1 activation by a variety of physiological signals. To test its role in response to feeding, L-TSC2-WT and -5A mice were fasted or fasted and refed for 1 hour. Plasma insulin was significantly elevated in refed compared to fasted mice, with no differences between L-TSC2-WT and -5A mice (**Figure S2D**). Somewhat surprisingly, mTORC1 signaling was induced by refeeding to a similar extent in both L-TSC2-WT and L-TSC2-5A livers (**Figure 2C**). Since hepatic mTORC1 is disconnected from insulin regulation in L-TSC2-5A mice, these results indicate that insulin signaling is not essential for mTORC1 activation by feeding in the liver. This result is consistent with fasting and refeeding studies in whole-body TSC2-WT and -5A mice, in which mTORC1 activation by feeding was retained in TSC2-5A livers, despite a clear defect in feeding induced activation of mTORC1 in skeletal muscle^48^. Together, these data support a model whereby nutrients or nutrient-derived signals independent of insulin are sufficient for hepatic mTORC1 activation in response to feeding a normal chow diet.

### Amino acids are necessary and sufficient for hepatic mTORC1 activation

Given that mTORC1 activation in cell culture models is dependent on amino acids^50^, we hypothesized that dietary protein content might override the insulin response in the activation of hepatic mTORC1 signaling by feeding. Indeed, quantitative metabolomics revealed that plasma concentrations of nearly all amino acids were elevated upon two hours of feeding relative to fasted mice, with concentrations of alanine, glutamine, valine, and proline being significantly increased with feeding in this analysis (**Figure 3A**). To test whether dietary protein is required for feeding to induce hepatic mTORC1 activation in L-TSC2-5A mice, mice were fasted and refed with a protein-free (PF) diet or an isocaloric semi-purified control diet (CTL) for one hour (**Figure 3B**). Relative to L-TSC2-WT livers, AKT activation was enhanced in the L-TSC2-5A livers under all conditions, but mTORC1 signaling was only significantly reduced in the CTL diet-induced L-TSC2-5A livers (**Figure 3C**, see graphs for quantification). The reduced mTORC1 activation in response to CTL diet refeeding in L-TSC2-5A livers is distinct from the response to chow diet refeeding (**Figure 2C**), which activated mTORC1 to a similar extent in L-TSC2-WT and -5A livers. Interestingly, mTORC1 activation was significantly blunted by refeeding the PF diet in both L-TSC2-WT and -5A livers (**Figure 3C**), despite significant increases in blood glucose (**Figure S3A**) and plasma insulin (**Figure 3D**) in both genotypes. Plasma insulin was lower in response to the PF diet relative to control diet, which likely reflects the established contribution of amino acids to pancreatic insulin secretion^51, 52^. However, insulin levels were still significantly induced over fasted (∼five-fold) by this diet and upstream activation of AKT was enhanced relative to CTL diet. These data demonstrate that dietary protein is essential for hepatic mTORC1 activation by feeding in a manner that is dominant over insulin signaling to AKT and TSC2.

**Figure 3.**
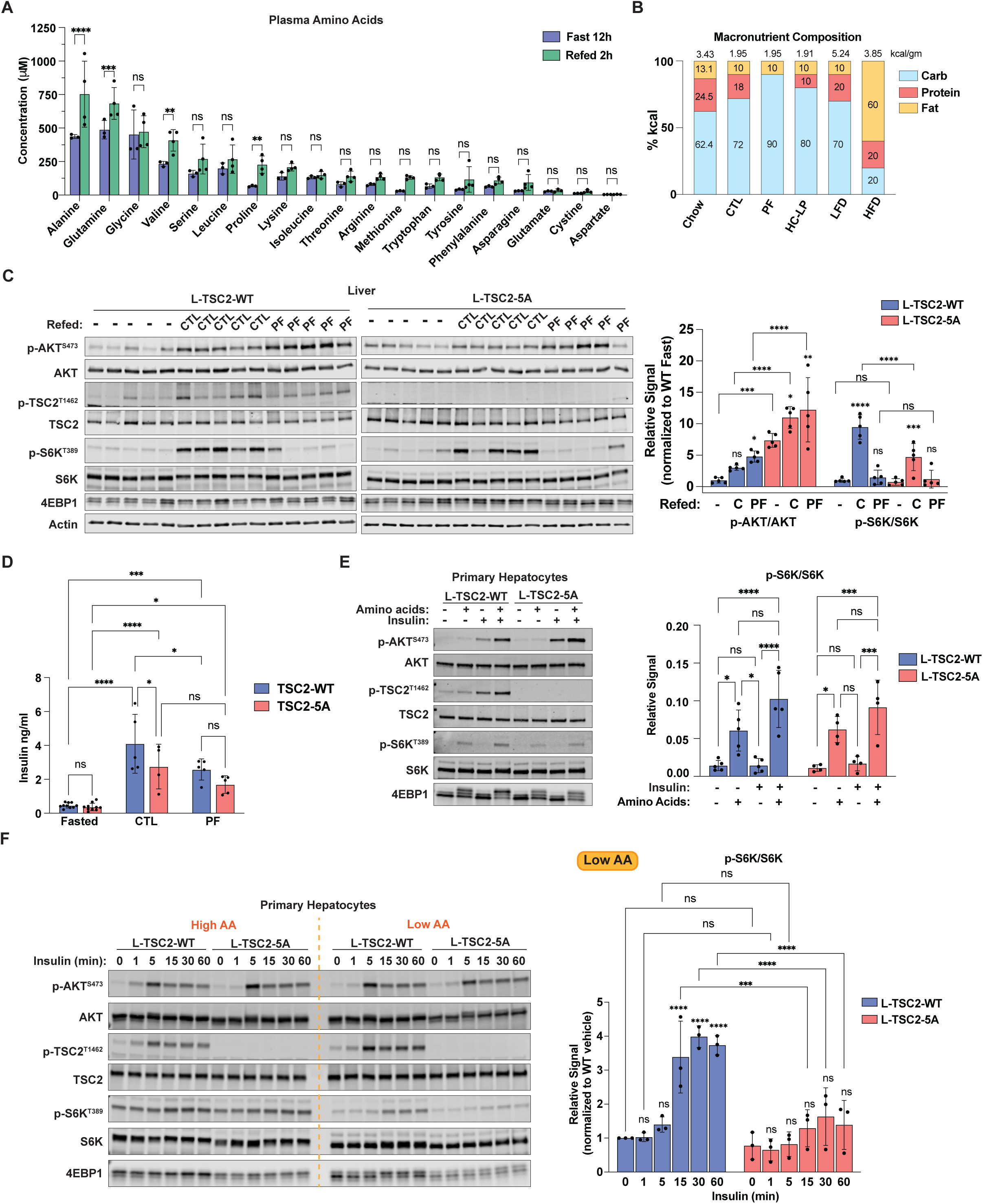
Amino acids are necessary and sufficient for hepatic mTORC1 activation. (A) Male TSC2-WT (whole-body) mice were fasted for 12 h (daytime) and, where indicated, refed for 2 h. The absolute concentrations of amino acids (histidine not detected) were analyzed in plasma from blood collected serially prior to and following refeeding and are graphed as mean ± SD; n = 3 fasted, 4 refed. (B) Graph depicting macronutrient composition of diets used throughout the study, with caloric density of diet (kcal/gram) depicted at top of each bar. (C) Immunoblots of liver lysates from 9-week-old L-TSC2-WT or -5A male mice fasted for 14 h overnight and, where indicated, refed for 1 h with isocaloric control (CTL) or protein-free (PF) diets, with accompanying bar graphs of the mean ± SD phospho-to-total protein signal quantifications normalized to the L-TSC2-WT fasted group; n = 5/group. The notations above each bar indicate significance relative to the respective fasted control group for each genotype. L-TSC2-WT and -5A lysates were analyzed on separate gels, but differences in gel loading were accounted for by normalizing to total protein signals for each target. (D) Plasma insulin from mice in (C) graphed as mean ± SD. Blood samples for refed mice were collected 30 min following refeeding; n = 5/group. (E) Immunoblot analysis of lysates from primary mouse hepatocytes isolated from L-TSC2-WT and -5A mice cultured overnight in serum-free media, deprived of amino acids for 1 h, then stimulated with amino acids or insulin (10 nM) alone or in combination for 15 min, with accompanying bar graphs of the mean ± SD of phospho-to-total S6K1 signal quantifications; n = 4-5 biological replicates. There was no significant difference in signal between L-TSC2-WT and -5A hepatocytes for any condition. (F) Immunoblot analysis of lysates from primary mouse hepatocytes isolated from L-TSC2-WT and -5A mice cultured overnight in serum-free DMEM (High AA) or DMEM containing 10% of its amino acid content (Low AA), followed by stimulation with insulin (10 nM) for the indicated times. The accompanying bar graphs of the mean ± SD phospho-to-total S6K1 signal quantifications for the low amino acid condition is depicted as fold induction from the L-TSC2-WT vehicle-treated (0 time point) condition (calculated separately for each replicate experiment). The notations directly above each bar indicate whether the signal was significantly induced from its respective vehicle control (0 time point) for each genotype; n = 3 biological replicates. Statistical analysis: (A) mixed-effects analysis with Fisher’s LSD test; (C-F) two-way ANOVA with Šidák correction. Not significant (ns = p≥0.05), *p < 0.05, **p < 0.01, ***p < 0.001, and ****p < 0.0001.

In most settings, full activation of mTORC1 is thought to require signals from both nutrients and exogenous growth factors or hormones^5^. However, the results above suggest that dietary nutrients are dominant over insulin signaling for the activation of mTORC1, at least in the liver. To test this hierarchy further, L-TSC2-WT and -5A primary mouse hepatocytes were depleted of amino acids for one hour followed by stimulation with amino acids and insulin alone or in combination for 15 minutes. Amino acids were sufficient to robustly activate mTORC1 in both L-TSC2-WT and -5A hepatocytes, while insulin alone did not stimulate mTORC1 in either genotype (**Figure 3E**). As expected, neither AKT activation nor TSC2 phosphorylation were induced by amino acids alone but were stimulated by insulin. The addition of insulin with amino acids modestly increased mTORC1 induction compared to amino acids alone, but this effect was not significant. These results indicate that amino acids are necessary and sufficient for near maximal mTORC1 activation in hepatocytes, providing an insulin-independent mechanism capable of activating hepatic mTORC1 in L-TSC2-5A mice upon feeding a normal chow diet.

To further determine the interaction between the insulin response and amino acid concentrations in the regulation of hepatic mTORC1 signaling, primary L-TSC2-WT and-5A hepatocytes were treated with a time course of insulin in the presence of high and low amino acids, while keeping all other media components constant. Insulin acutely stimulated AKT activation and TSC2 phosphorylation similarly in high and low amino acid media (**Figure 3F**). However, basal mTORC1 signaling was greatly elevated in the high amino acid media, with only modest further induction by insulin (**Figure 3F, S3B**). Along with lower basal mTORC1 activity, mTORC1 signaling was more overtly stimulated by insulin under low amino acids conditions in L-TSC2-WT hepatocytes, and this induction was almost entirely abolished in L-TSC2-5A hepatocytes (**Figure 3F**). These results indicate that amino acid levels dictate basal hepatic mTORC1 activation and its dynamic sensitivity to insulin stimulation, with insulin signaling to mTORC1 being dependent on AKT-mediated phosphorylation of TSC2.

### Dietary carbohydrate-to-protein ratio governs the contribution of insulin-AKT-TSC2 signaling to hepatic mTORC1 activation

The results above provide evidence that amino acid availability determines the role of insulin signaling in the activation of hepatic mTORC1. It is interesting to note that synthetic CTL diet control for the PF diet has a higher proportion of carbohydrates to protein than the chow diet (**Figure 3B**), and the L-TSC2-5A mice showed a significant reduction in the stimulation of hepatic mTORC1 signaling by refeeding this diet, relative to their L-TSC2-WT counterparts (**Figure 3C**). To further test whether the dietary carbohydrate-to-protein ratio dictates the dominant route to mTORC1 activation in liver tissue, the response of L-TSC2-WT and -5A mice to refeeding a high carbohydrate, low protein (HC-LP) diet was determined. A semi-purified diet containing 80% calories from carbohydrate and 10% from protein was formulated for this purpose (**Figure 3B**), as previous studies have demonstrated that short-term feeding of a 10% protein diet does not elicit markers of protein restriction^53^. Following one-hour refeeding with the HC-LP diet, hepatic mTORC1 signaling was induced by approximately seven-fold in L-TSC2-WT mice, but induction was only modest and significantly reduced in L-TSC-5A livers (**Figure 4A**). These results demonstrate that lower dietary protein content renders AKT-mediated phosphorylation of TSC2 the primary route to feeding-induced hepatic mTORC1 signaling.

**Figure 4.**
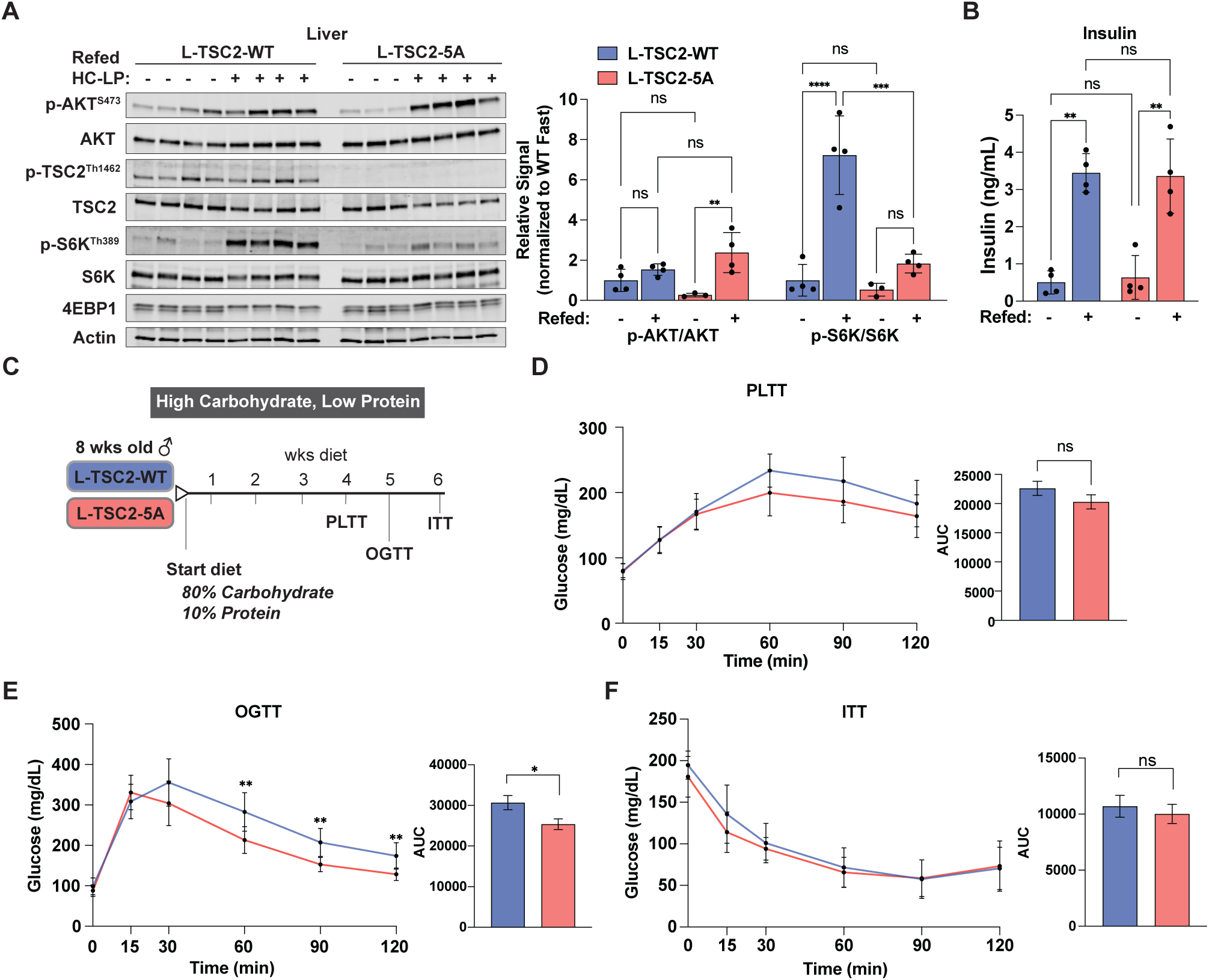
Dietary protein-to-carbohydrate ratio governs the contribution of AKT-TSC2 signaling to hepatic mTORC1 activation. (A) Immunoblots of liver lysates from 12-week-old L-TSC2-WT or -5A male mice fasted for 14 h overnight and, where indicated, refed for 1 h with a high carbohydrate, low protein (HC-LP) diet, with accompanying bar graphs the mean ± SD phospho-to-total protein signal quantifications normalized to the L-TSC2-WT fasted group; n = 3-4/group. (B) Plasma insulin of mice from (A) graphed as mean ± SD. Tail vein blood samples were collected from fasted mice and 30 minutes after refeeding; n = 4 /group. (C-F) Male L-TSC2-WT or -5A mice were fed HC-LP diet starting at eight weeks of age. (C) Timeline of diet study and phenotypic characterizations. (D) Pyruvate-lactate tolerance test performed after four weeks diet. Mice were fasted for 16 h overnight followed by i.p. injection with lactate (1.5 mg/kg) and pyruvate (150μg/kg). Blood glucose measurements over time are plotted as mean ± SD. AUC values are graphed as mean ± SEM; n = 11 WT, 12 5A. (E) Oral glucose tolerance test performed after five weeks diet. Mice were fasted for 16 h overnight followed by oral gavage of glucose (2 g/kg). Blood glucose measurements over time are plotted as mean ± SD. AUC values are graphed as mean ± SEM; n = 11 WT, 10 5A. (F) Insulin tolerance test performed after six weeks diet. Mice were fasted for 2 h followed by i.p. injection with insulin (0.5 U/kg). Blood glucose measurements over time are plotted as mean ± SD. AUC values are graphed as mean ± SEM; n = 8 WT, 9 5A. Statistical analysis: (A and B) two-way ANOVA with Šidák correction; (D-F) two-way ANOVA with Šidák correction for curves, Welch’s t-test for AUC. Not significant (ns = p≥0.05), *p < 0.05, **p < 0.01, ***p < 0.001, and ****p < 0.0001.

While there are no known direct effects of mTORC1 signaling on hepatic or systemic glucose metabolism, mTORC1-mediated feedback mechanisms can influence the insulin responsiveness of a given tissue^7, 8, 28, 54–57^. Importantly, plasma insulin measured in samples taken 30 minutes after refeeding the HC-LP diet were similar between L-TSC2-WT and -5A mice (**Figure 4B**), but the induction of AKT activation was more pronounced in the L-TSC2-5A livers, suggesting enhanced hepatic insulin sensitivity (**Figure 4A**). To test effects on glucose homeostasis, we assayed pyruvate, glucose and insulin tolerance in L-TSC2-WT and -5A mice during prolonged HC-LP diet feeding (**Figure 4C**). As on a normal chow diet, no differences in body weight, lean mass, or fat mass were observed between L-TSC2-WT and -5A mice fed this diet for six weeks (**Figures S3C-E**). Additionally, fasting plasma insulin was not significantly different between L-TSC2-WT and -5A mice (**Figure S3F**). After four weeks on the HC-LP diet, a pyruvate-lactate tolerance test was performed on fasted L-TSC2-WT and -5A mice to measure gluconeogenic capacity. While there was a trend towards reduced endogenous glucose production in L-TSC2-5A compared to L-TSC2-WT mice, the glucose values were not significantly different (**Figure 4D**). In response to an oral glucose tolerance test performed after five weeks on the HC-LP diet, L-TSC2-5A mice exhibited a modest, but significant, improvement in glucose clearance (**Figure 4E**). However, systemic insulin sensitivity was not significantly different between L-TSC2-WT and -5A mice after six weeks on this diet (**Figure 4F**). Together, these data demonstrate that AKT-mediated TSC2 phosphorylation regulates mTORC1 activation in the context of a low protein diet, which likely through feedback regulation of the hepatic insulin response exerts moderate effects on systemic glucose homeostasis.

### Hyperinsulinemia accompanying diet-induced obesity is not associated with chronic activation of hepatic mTORC1 signaling

Next, we used a parallel approach to evaluate hepatic mTORC1 activation in response to the pathological states of obesity and hyperinsulinemia. C57BL/6J male mice were fed a high fat diet (HFD) containing 60% kilocalories from fat or a low-fat control diet (LFD) containing 10% fat beginning at eight weeks of age. Dietary protein was constant between the HFD and LFD. Systemic glucose homeostasis was evaluated by glucose and insulin tolerance tests in cohorts fed HFD or LFD diets for 4, 8, or 16 weeks (**Figure 5A**). mTORC1 signaling was measured in liver tissue collected from these same mice following a daytime fast (**Figure 5A**). As expected, HFD-fed mice weighed significantly more than LFD-fed mice starting after just one week of diet (**Figure 5B**). This elevated body weight was associated with increased fat mass in HFD versus LFD mice after 4 and 8 weeks of diet (**Figure S4A**), as well as significantly higher lean mass at 8 weeks HFD (**Figure S4B**). Plasma insulin levels were also elevated in HFD compared to LFD mice at 4, 8, and 15 weeks, and were significantly higher when measured at 8 and 15 weeks (**Figure 5C**). At 4 weeks, systemic insulin sensitivity was significantly impaired in HFD compared to LFD mice (**Figure 5D**), with progressive impairments at 8 and 16 weeks (**Figures 5F, H**). Glucose tolerance was similarly impaired in HFD-fed mice (**Figure S4C, D**). Strikingly, mTORC1 signaling was not significantly elevated in the liver tissue of HFD compared to LFD mice, even after 16 weeks, despite the overt obesity, hyperinsulinemia, and insulin resistance phenotypes of these mice (**Figure 5E, G, I**). Accompanying the chronic hyperinsulinemia was significantly elevated hepatic Akt activation in the HFD compared to LFD mice at 16 weeks, while total IRS1 protein levels, an indicator of insulin sensitivity, were reduced in these livers (**Figure 5I**). Collectively, these data are in opposition to the purported dogma that hepatic mTORC1 is chronically elevated in diet-induced obesity and contributes to systemic insulin resistance. However, they are in alignment with our results in L-TSC2-5A mice demonstrating that insulin is not the predominant signal driving mTORC1 activation in the liver.

**Figure 5.**
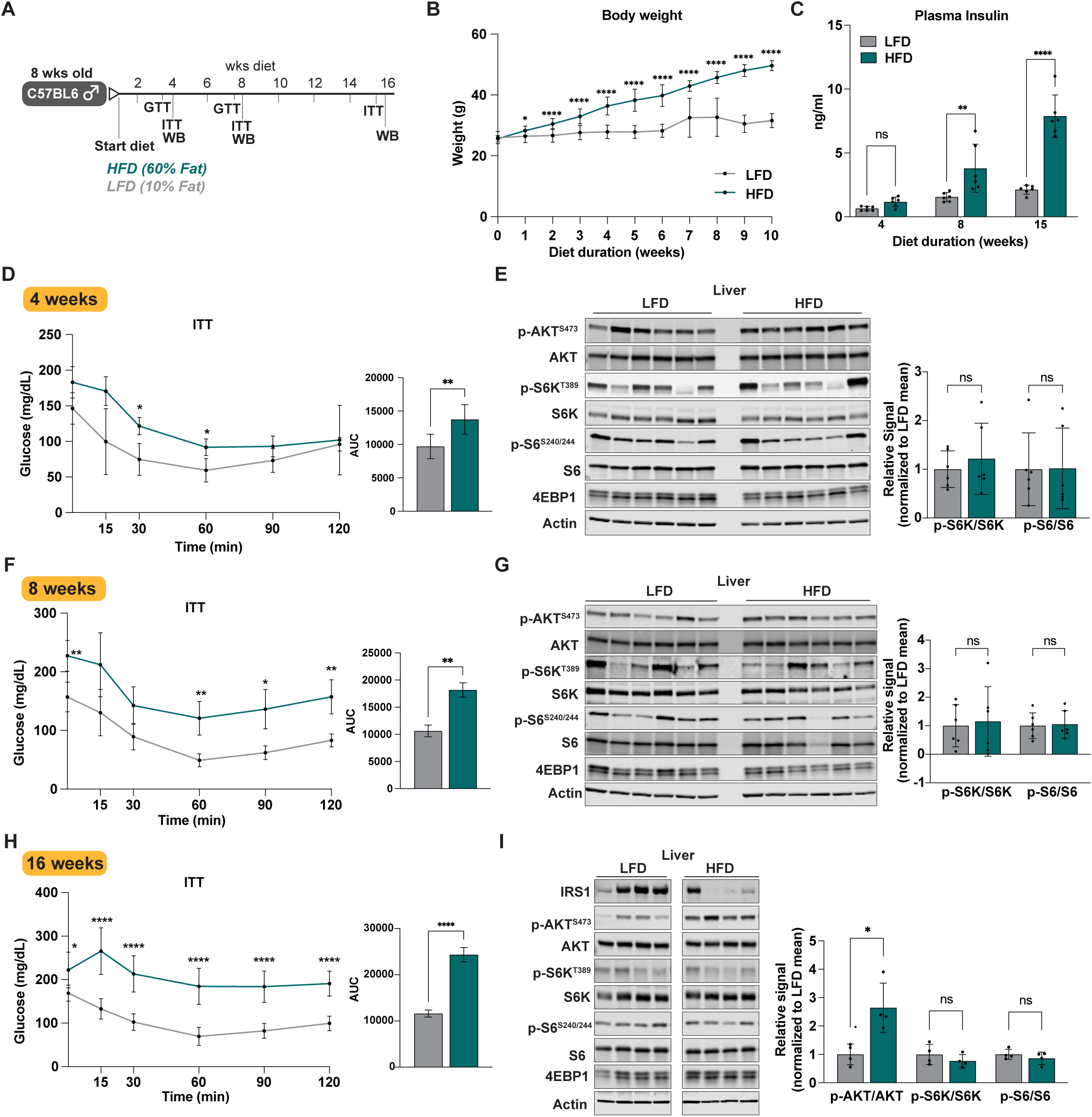
Hyperinsulinemia accompanying diet-induced obesity is not associated with chronic activation of hepatic mTORC1 signaling. (A-I) Male C57BL/6J mice were fed a high-fat diet (HFD) or low-fat control diet (LFD) beginning at eight weeks of age. (A) Timeline of diet study and phenotypic characterizations. (B) Body weight measurements of male C57BL/6J mice fed high fat or low-fat control diets graphed over time as mean ± SD; n = 6-18/group/timepoint. (C) Plasma insulin from 6-h fasted mice fed HFD or LFD diets for 4, 8, or 15 weeks graphed as mean ± SD; n = 6/group. (D-F) Insulin tolerance test performed after 3.5 (D), 7.5 (E), and 13.5 (F) weeks diet. Mice were fasted for 6 h followed by i.p. injection with insulin (0.5 U/kg). Blood glucose measurements over time are plotted as mean ± SD. AUC values are graphed as mean ± SEM; n = 6/group. (E) Immunoblots of liver lysates from mice in (D) fed HFD or LFD for 4 weeks and fasted for 6 h prior to tissue collection. Accompanying bar graphs of mean ± SD phospho-to-total protein signal quantifications normalized to the LFD group are shown; n = 6/group. (F) Insulin tolerance test performed after 7.5 weeks diet. Mice were fasted for 6 h followed by i.p. injection with insulin (0.5 U/kg). Blood glucose measurements over time are plotted as mean ± SD. AUC values are graphed as mean ± SEM; n = 5 LFD, 6 HFD. (G) Immunoblots of liver lysates from mice in (F) fed HFD or LFD for 8 weeks and fasted for 6 h prior to tissue collection. Accompanying bar graphs of mean ± SD phospho-to-total protein signal quantifications normalized to the LFD group are shown; n = 6/group. (H) Insulin tolerance test performed after 13.5 weeks diet. Mice were fasted for 6 h followed by i.p. injection with insulin (0.5 U/kg). Blood glucose measurements over time are plotted as mean ± SD. AUC values are graphed as mean ± SEM; n = 11 LFD, 10 HFD. (I) Immunoblots of liver lysates from mice in (H) fed HFD or LFD for 16 weeks and fasted for 12 h (daytime) prior to tissue collection. Accompanying bar graphs of mean ± SD phospho-to-total protein signal quantifications normalized to the LFD group are shown; n = 4/group. Statistical analysis: (B) mixed-effects analysis with Šidák’s correction; (C) two-way ANOVA with Šidák correction; (D) mixed-effects analysis with Šidák correction for curves, Welch’s t-test for AUC; (E, G, and I) Mann-Whitney test; (F and H) two-way ANOVA with Šidák correction for curves, Welch’s t-test for AUC. Not significant (ns = p≥0.05), *p < 0.05, **p < 0.01, ***p < 0.001, and ****p < 0.0001.

We previously demonstrated that, in contrast to the liver, muscle mTORC1 activation in response to feeding is largely dependent on AKT-mediated TSC2 phosphorylation^48^, suggesting that insulin may play a more dominant regulatory role for mTORC1 in muscle tissue. In agreement with this, activation of mTORC1 signaling was higher in gastrocnemius muscle of HFD-versus LFD-fed mice after 15 weeks (**Figure S4E**). These data highlight that different regulatory mechanisms contribute to mTORC1 activation across metabolic tissues. Whether the elevated mTORC1 activation in muscle of HFD-fed mice contributes to systemic insulin resistance or impaired glucose clearance requires further study.

### Obese L-TSC2-5A mice are not protected from metabolic impairment but maintain defects in insulin-stimulated hepatic mTORC1

We next evaluated whether chronic hyperinsulinemia driven by diet-induced obesity could accentuate differences between L-TSC2-WT and -5A mice, as seen with the HC-LP diet. We hypothesized that if hyperinsulinemia contributes to the pathological activation of hepatic mTORC1 it would be lower in L-TSC2-5A mice following HFD feeding and potentially accompanied by improvements in glucose homeostasis. To test this, L-TSC2-WT and -5A male and female mice were placed on a high-fat diet (HFD) at 8 weeks of age, with glucose and insulin tolerance tests performed after 8-10 weeks and again after 15-16 weeks of diet (**Figure 6A**). There were no significant differences in body weight or body composition between L-TSC2-WT and -5A mice across high fat diet feeding (**Figure 6B-D** (males), **Figure S5D-F** (females)). Plasma insulin levels increased progressively and to a similar extent in both L-TSC2-WT and -5A mice during HFD feeding (**Figure S5A** (males), **Figure S5G** (females)). Systemic glucose tolerance and insulin sensitivity were not significantly different between L-TSC2-WT and -5A mice after 8-10 weeks (data not shown) or 16 weeks HFD (**Figure 6E-F** (males), **Figure S5H-I** (females)). Additionally, hepatic glycogen levels were similar between L-TSC2-WT and -5A mice (**Figure S5B**). Plasma albumin, a marker of liver function, was also similar between L-TSC2-WT and -5A mice fed HFD (**Figure S5C**). These data indicate that L-TSC2-5A mice are not protected from systemic metabolic impairments association with diet-induced obesity.

**Figure 6.**
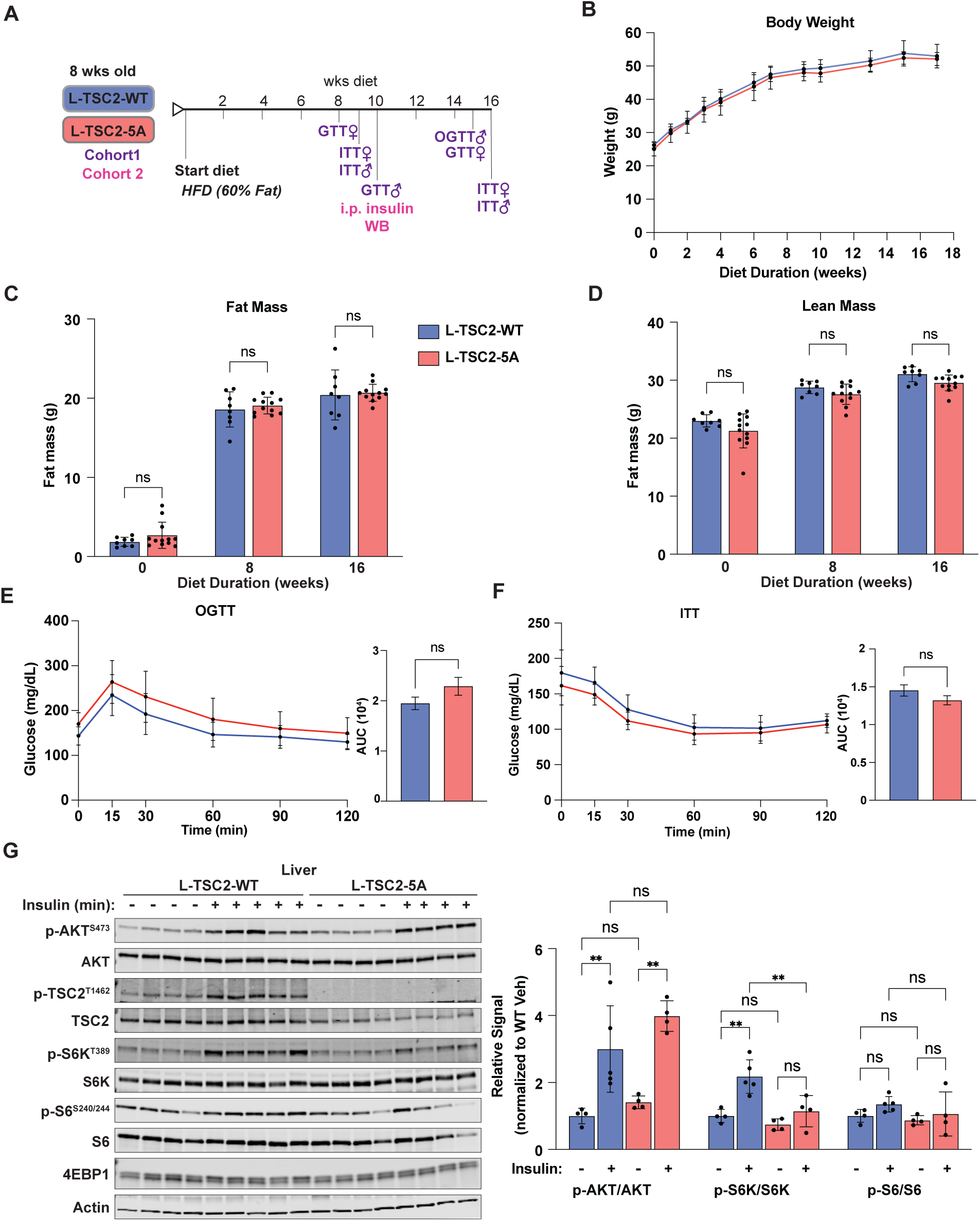
Obese L-TSC2-5A mice are not protected from metabolic impairment but maintain defects in insulin-stimulated hepatic mTORC1. (A-G) L-TSC2-WT and -5A male and female mice were fed high-fat diet (HFD) beginning at 8 weeks of age. (A) Schematic of HFD study in L-TSC2-WT and-5A mice with timeline of phenotypic characterizations. (B) Body weight measurements of L-TSC2-WT and -5A male mice fed HFD for indicated amount of time graphed as mean ± SD; n = 8 WT, 12 5A. (C, D) Fat (C) and lean (D) mass of L-TSC2-WT and -5A male mice at baseline (0 weeks on diet; eight weeks of age) and after 8 or 16 weeks HFD, measured by EchoMRI and graphed as mean ± SD; n = 8 WT, 12 5A. (E) Oral glucose tolerance test performed after 15 weeks HFD. Mice were fasted for 6 h followed by oral gavage of glucose (1 g/kg). Blood glucose measurements over time are plotted as mean ± SD. AUC values are graphed as mean ± SEM; n = 8 WT, 12 5A. (F) Insulin tolerance test performed after 16 weeks HFD. Mice were fasted for 6 h followed by i.p. injection with insulin (0.5 U/kg). Blood glucose measurements over time are plotted as mean ± SD. AUC values are graphed as mean ± SEM; n = 8 WT, 11 5A. (G) Immunoblots of liver lysates from L-TSC2-WT or -5A mice fed HFD for 10 weeks and fasted for 14 h overnight then injected i.p. with vehicle (saline) or insulin (0.5 U/kg) for 20 min. Accompanying bar graphs of mean ± SD phospho-to-total protein signal quantifications normalized to the L-TSC2-WT vehicle-treated group are shown; n = 4-5/group. Statistical analysis: (B-D and G) two-way ANOVA with Šidák’s correction; (E) mixed-effects analysis with Šidák correction for curves, Welch’s t-test for AUC; (F) two-way ANOVA with Šidák correction for curves, Welch’s t-test for AUC. Not significant (ns = p≥0.05), *p < 0.05, **p < 0.01, ***p < 0.001, and ****p < 0.0001.

To test whether fasted or insulin-stimulated hepatic mTORC1 activation was altered in L-TSC2-5A compared to - WT mice under conditions of obesity and hyperinsulinemia, we assessed liver signaling in fasted L-TSC2-WT and -5A males treated with vehicle or insulin after 10 weeks HFD. Fasting hepatic mTORC1 activation was not significantly different between L-TSC2-WT and -5A mice (**Figure 6G**). Importantly, as detected in lean mice (**Figure 2A**), mTORC1 induction by exogenous administration of insulin was impaired in diet-induced obese L-TSC2-5A compared to - WT mice, but this was not associated with significantly enhanced AKT activation (**Figure 6G**). These data further support that hyperinsulinemia is not a primary driver of hepatic mTORC1 activation.

### Glucagon is a suppressive signal of hepatic mTORC1

The present study has focused on the interplay between insulin and nutrients in the induction of hepatic mTORC1 upon feeding. However, whether there are specific signals that suppress hepatic mTORC1 activation during fasting, outside of a reduction in feeding-induced stimulatory signals, is unknown. To this end, the fasting hormone glucagon has been reported to inhibit mTORC1 in perfused rat liver and primary hepatocytes^58–60^, and forskolin, which recapitulates one aspect of glucagon signaling by increasing cAMP levels and activating protein kinase A (PKA), also suppresses mTORC1 in non-hepatic cell lines^59, 61^. However, the potential contribution of glucagon to the suppression of hepatic mTORC1 during fasting has not been directly tested.

In a time course experiment, we found that glucagon acutely suppresses mTORC1 signaling in primary mouse hepatocytes, with maximal inhibition at 30 min and sustained inhibition for at least 6 hours (**Figure 7A**). The induction of canonical glucagon signaling was confirmed by the PKA-mediated phosphorylation of CREB. Acute glucagon treatment was capable of attenuating insulin-induced mTORC1 activation in primary hepatocytes (**Figure 7B**). Glucagon also inhibited the elevated mTORC1 signaling of hepatocytes with siRNA-mediated knockdown of *Tsc2*, suggesting a TSC complex-independent mechanism of suppression (**Figure 7C**).

**Figure 7.**
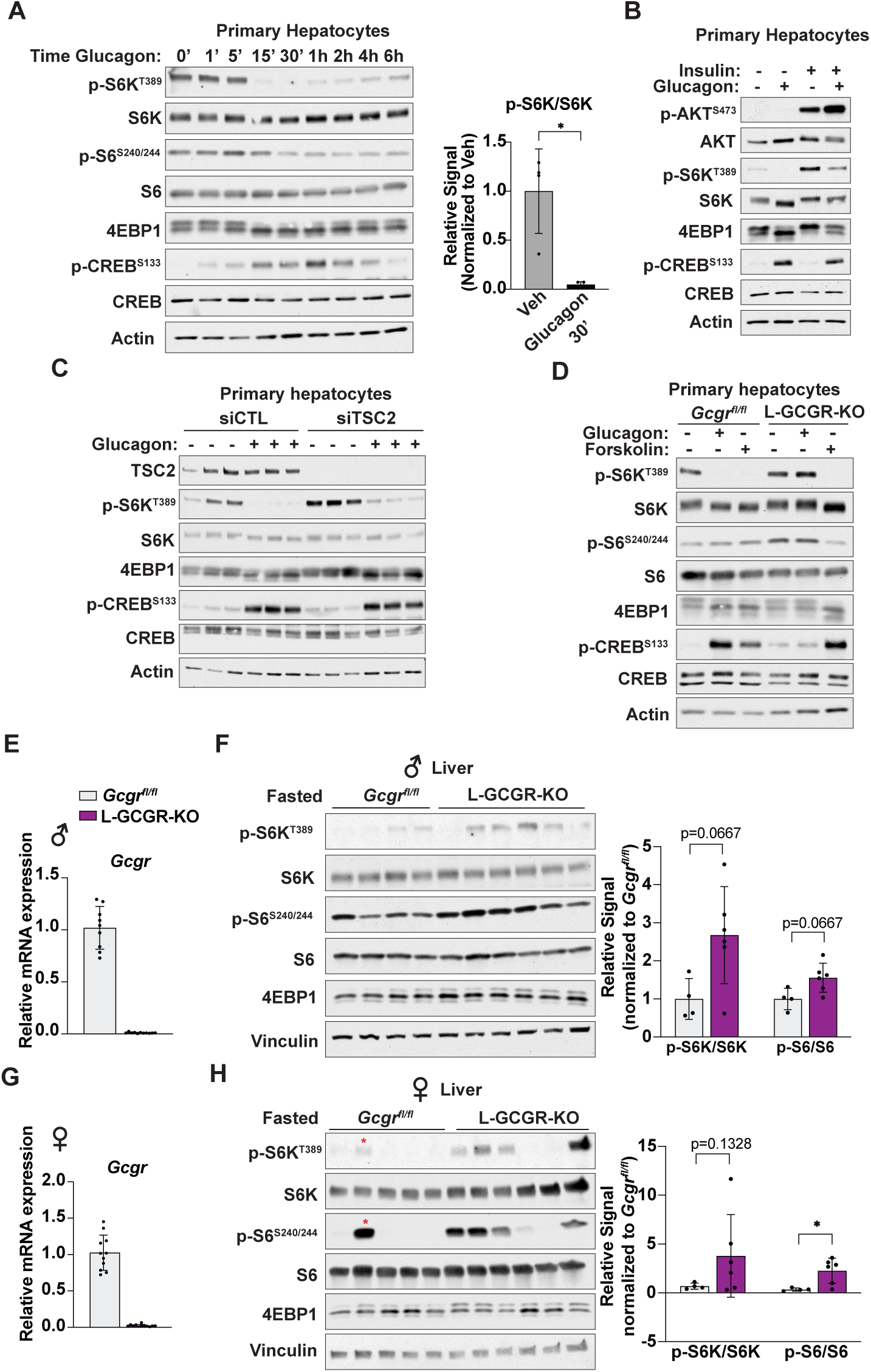
Glucagon is a suppressive signal of hepatic mTORC1. (A) Immunoblots of lysates from C57BL/6J primary mouse hepatocytes that were serum-starved overnight then treated with glucagon (10 nM) for the indicated times. Accompanying bar graph of the mean ± SD phospho-to-total S6K1 signal quantifications for the vehicle (0 time point) and 30 min glucagon treatment normalized to the vehicle-treated group are shown; n = 4 biological replicates. (B) Immunoblots of lysates from C57BL/6J primary mouse hepatocytes that were serum starved overnight then pre-treated with insulin (100 nM) for 30 min followed by treatment with glucagon (10 nM) for 30 min, where indicated. (C) Immunoblots of lysates from C57BL/6J primary mouse hepatocytes with siRNA-mediated knockdown *Tsc2* or with a scrambled control siRNA (siCTL) for 72 h followed by overnight serum deprivation and treatment with glucagon (10 nM) for 30 min; n = 3 technical replicates. (D) Immunoblots of lysates from *Gcgr^fl/fl^* or L-GCGR-KO primary mouse hepatocytes serum-starved overnight and treated with glucagon (10 nM) or forskolin (10 μM) for 30 min, where indicated. (E) RT-qPCR quantification of *Gcgr* transcripts in liver tissue of male *Gcgr^fl/fl^* and *L-Gcgr-KO* mice graphed as mean ± SD relative to *Gcgr^fl/fl^*; n = 9 *Gcgr^fl/fl^*, 12 *L-Gcgr-KO*. (F) Immunoblots of liver lysates from male *Gcgr^fl/fl^* and *L-Gcgr-KO* mice fasted for 12 h (daytime) prior to tissue collection. Accompanying bar graphs of mean ± SD phospho-to-total protein signal quantifications normalized to the *Gcgr^fl/fl^* group are shown; n = 4 *Gcgr^fl/fl^*, *6 L-Gcgr-KO*. (G) RT-qPCR quantification of *Gcgr* transcripts in liver tissue of female *Gcgr^fl/fl^* and *L-Gcgr-KO* mice graphed as mean ± SD relative to *Gcgr^fl/fl^*; n = 11 *Gcgr^fl/fl^*, 12 *L-Gcgr-KO*. (H) Immunoblots of liver lysates from female *Gcgr^fl/fl^* and *L-Gcgr-KO* mice fasted for 12 h (daytime) prior to tissue collection. Accompanying bar graphs of mean ± SD phospho-to-total protein signal quantifications normalized to the *Gcgr^fl/fl^*group are shown; n = 4 *Gcgr^fl/fl^*, *6 L-Gcgr-KO*. Red asterisks indicate a sample that was removed from analysis for both S6K1 and S6 signal quantification, as it was found to be a significant outlier by the Grubb’s test. Statistical analysis: (A, F, and H) Mann-Whitney test. Not significant (ns = p≥0.05), *p < 0.05, **p < 0.01, ***p < 0.001, and ****p < 0.0001.

We next took a genetic approach to evaluate the role of glucagon signaling in the regulation of hepatic mTORC1. *Gcgr^fl/fl^* mice were bred with mice expressing Albumin-Cre to generate mice with liver-specific knockout of the glucagon receptor (L-GCGR-KO), as previously described^62^. We first assessed whether the glucagon receptor (GCGR) is required for the glucagon-mediated suppression of mTORC1 using hepatocytes from L-GCGR-KO or *Gcgr^fl/fl^* mice. Glucagon treatment had no effect on mTORC1 signaling in *Gcgr*^-/-^ hepatocytes, while forskolin, which activates the cAMP signaling cascade downstream of GCGR, suppressed mTORC1 signaling similarly in *Gcgr^fl/fl^* and L-GCGR-KO hepatocytes, with corresponding effects on CREB phosphorylation (**Figure 7D**). These data indicate that suppression of mTORC1 by glucagon is mediated through the hepatic GCGR.

To assess whether hepatic glucagon signaling is required for the suppression of hepatic mTORC1 during fasting *in vivo*, mTORC1 signaling was analyzed in liver tissue of fasted *Gcgr^fl/fl^*and L-GCGR-KO mice. Loss of *Gcgr* expression was verified in liver tissue by RT-qPCR (**Figures 7E, G**). Phosphorylation of markers of mTORC1 signaling were modestly elevated in the livers of both male and female L-GCGR-KO mice compared to *Gcgr^fl/fl^* mice (**Figure 7F, 7H**). Together, these data provide support that glucagon can inhibit hepatic mTORC1 signaling and may contribute to its suppression in the fasted state.

## Discussion

The liver coordinates systemic metabolism in response to physiological nutrient and energy supply. The mTORC1 signaling network controls cellular metabolism and is sensitive to local and systemic nutrient status but physiological regulatory mechanisms in the liver are poorly defined. By mutating TSC2 at the five AKT-mediated phosphosites that are required for mTORC1 activation by insulin in cells^24, 26, 48^, the L-TSC2-5A mouse model serves as a genetic tool to define the specific contribution of insulin-AKT-TSC signaling to the physiological regulation of mTORC1. This approach is distinct from previously published mouse models, which are characterized by genetic loss-of-function of one of the nodes in the signaling network. These previous studies demonstrate that the TSC complex is required for the dynamic regulation of hepatic mTORC1 by fasting and feeding^7, 8, 28^, but the specific role of the AKT-TSC2 circuit has not been defined.

Studies on AKT1/2 knockout mice demonstrate that AKT is required for hepatic mTORC1 activation in response to insulin, but metabolic perturbations in these mice, including hyperglycemia and hyperinsulinemia, may indirectly influence mTORC1 regulation^33, 35^. The present study establishes that Akt-mediated TSC2 phosphorylation is the primary mechanism by which insulin and systemic glucose activate mTORC1 in the liver. However, this does not rule out the potential contribution of alternative AKT-dependent mechanisms influencing other mTORC1 suppressors, such as PRAS40^14, 15, 63^ or AMPK^17, 64^, to the residual activation of mTORC1 observed in the L-TSC2-5A liver by insulin or glucose. It is also possible that additional, unidentified AKT-mediated phosphorylation sites on the TSC complex sensitive to feeding, glucose, or insulin could contribute to hepatic mTORC1 induction. Furthermore, cellular glucose has been reported to activate mTORC1 in other settings via both Rag- and TSC complex-mediated mechanisms^40, 65–69^, as well as through the glycolytic intermediate dihydroxyacetone phosphate (or DHAP)^70^. It seems likely that different mechanisms mediate mTORC1 regulation by systemic and cellular glucose and depend on the tissue, duration, or concentration of glucose.

Our data demonstrate that dietary protein, but not insulin signaling, is essential for hepatic mTORC1 induction by feeding. Studies on mice expressing constitutively active RagA or lacking the leucine sensors Sestrin1 and Sestrin2 have provided genetic evidence that the amino acid sensing branch plays a critical role in hepatic mTORC1 regulation^40, 71–73^. However, these studies do not discern the relative influence of dietary nutrients and insulin signaling in hepatic mTORC1 regulation across the transition between fasting and feeding. Our findings indicate that amino acids are both necessary and sufficient for cell-intrinsic mTORC1 activation in hepatocytes and that amino acid levels dictate the importance of insulin-stimulated AKT-mediated phosphorylation of TSC2 for full mTORC1 activation in hepatocytes. Similarly, the essentiality of insulin signaling to postprandial hepatic mTORC1 activation was positively correlated with the dietary carbohydrate-to-protein ratio. These findings bolster the conclusion that nutrients are the primary signal in the regulation of hepatic mTORC1 (**Figure 8**), in contrast to many other settings where maximal mTORC1 activation requires both nutrients and insulin^5^. Indeed, previous work in whole-body TSC2-WT and -5A mice found that feeding-induced mTORC1 activation in muscle tissue is dependent on AKT-mediated TSC2 phosphorylation^48^. Thus, it will be important to further determine how established upstream pathways differentially impact mTORC1 in distinct tissue contexts.

**Figure 8.**
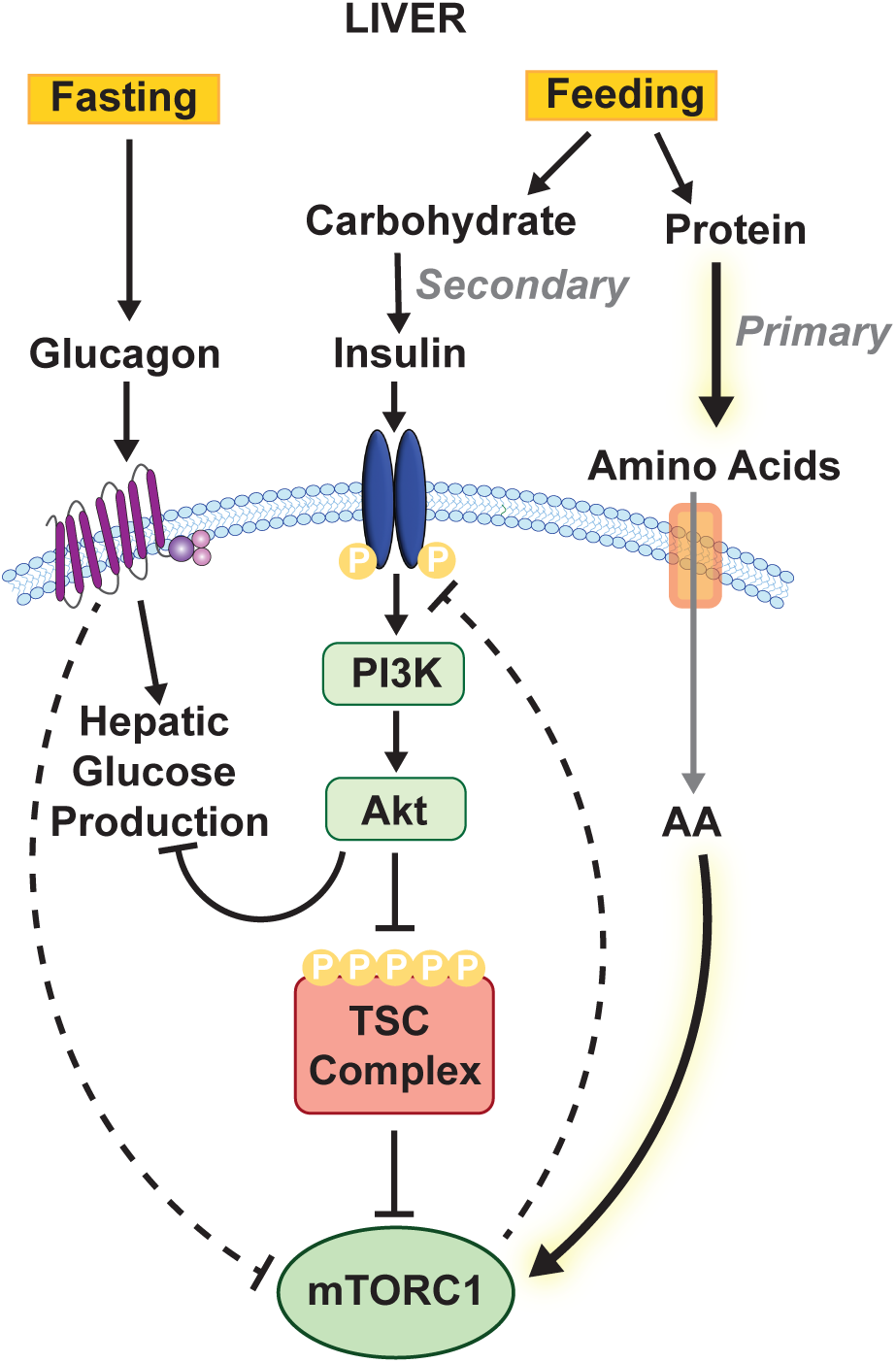
**Model of physiological regulation of hepatic mTORC1 by fasting and feeding.**

Despite our observations that oral delivery of glucose activates mTORC1 signaling in the liver, we found that refeeding with a protein-free diet containing 90% kilocalories from carbohydrate (48% sucrose) did not induce hepatic mTORC1 signaling. This discrepancy may be due to differences in timing (25 minutes glucose gavage versus one-hour feeding) and/or differences in nutrient absorption between glucose gavage and digestion following feeding. Additionally, dietary sucrose, which is metabolized into glucose and fructose may be less potent for mTORC1 induction than a bolus of pure glucose. Indeed, fructose was found to acutely suppress hepatic mTORC1 in mice^74^. It is also possible that gut- or nervous system-derived signals differentially activated by voluntary eating or digestion versus absorption of purified glucose contribute to the hepatic mTORC1 signaling response. To this end, POMC neuron-mediated activation of the sympathetic nervous system reportedly induces hepatic mTORC1 in response to food perception in mice^75^.

Published studies suggest that the dynamic nature of mTORC1 signaling in insulin responsive tissues, including the liver, is critical to whole-body metabolic homeostasis^9, 28, 37^. It has been proposed that liver mTORC1 activation becomes elevated in obesity and contributes to hepatic insulin resistance, with in vitro studies implementing negative feedback mechanisms on the IRS proteins and the insulin receptor^10, 43–45^. However, the mechanisms underlying aberrant hepatic mTORC1 activation under states of nutrient overload and obesity have not been directly tested. Our study found no differences in hepatic mTORC1 activation between mice fed a high fat versus low fat diet. Consistent with the lack of mTORC1 upregulation in response to hyperinsulinemia and hyperglycemia in diet-induced obesity, insulin-stimulated hepatic mTORC1 signaling had minimal contributions to impaired glucose homeostasis in obese L-TSC2-WT and -5A mice. Our findings contradict the purported dogma that hepatic mTORC1 is chronically upregulated in diet-induced obesity^10, 11, 76^, emphasizing the need for further investigation into the causal role of mTORC1 in hepatic insulin resistance and metabolic dysfunction. One potential explanation for the discrepancy is that some previous studies utilized a chow diet to control for HFD^9, 10^. Chow diets contain minimal sucrose and various whole food ingredients that semi-purified diets do not, confounding the comparison between obese and lean mice in these studies. Interestingly, obesity in humans is associated with elevated levels of branched-chain amino acids (BCAAs)^77^, which have been proposed to contribute to insulin resistance, at least in part, through liver and/or muscle mTORC1 activation^77–79^. Given our findings that dietary protein and amino acids are dominant for the activation of hepatic mTORC1, it is possible that a BCAA-mTORC1 axis drives metabolic impairments in obesity without the necessity of broad, persistent upregulation. Another possibility is that zonal regulation of hepatic mTORC1, which is not reflected in immunoblots of whole liver lysates, is impaired under conditions of obesity^72, 80, 81^. Identifying downstream targets of mTORC1 in the liver that mediate its feedback effects on insulin signaling or control of other metabolic pathways will increase our understanding of mTORC1 function in both physiology and metabolic disease.

In addition to investigating mechanisms of hepatic mTORC1 activation, we also evaluated glucagon, which inhibits mTORC1 in hepatocytes and perfused liver tissue^58–60^, as a potential mediator of hepatic mTORC1 suppression during fasting. We provide genetic evidence in support of hepatic mTORC1 regulation by glucagon *in vivo*. The mechanism of mTORC1 regulation by glucagon remains unclear but is dependent on the canonical glucagon receptor and appears to rely on cAMP signaling, as it is recapitulated by forskolin, and seems at least partly independent of the TSC complex^59, 61^. Glucagon promotes amino acid catabolism in hepatocytes and liver tissue^82^, which may contribute to its inhibition of mTORC1. However, it is likely that several mechanisms contribute to mTORC1 suppression during fasting, including a decrease in stimulatory signals from circulating amino acids and insulin. The metabolic response to fasting, including upregulation of glycogenolysis, gluconeogenesis, fatty acid oxidation, and ketogenesis, is impaired in L-GCGR-KO mice ^83–86^. Given that several of these processes are opposed by mTORC1^7, 8, 71^, it is possible that aberrant mTORC1 signaling contributes to the impaired fasting response in L-GCGR-KO mice, but this remains to be directly tested. Importantly, the regulation of glucagon secretion is defective in obese mice^87^, leading to diminished dynamic shifts in plasma ratios of glucagon to insulin, which could also contribute to aberrant mTORC1 regulation in metabolic disease.

This study demonstrates that macronutrient composition influences hepatic mTORC1 activation, independent of insulin induction, which could have important implications for energy balance and metabolic health. Understanding how mTORC1 is regulated and functions physiologically within each tissue in response to nutrient and hormonal cues will clarify its role in mediating the metabolic and anti-aging effects of dietary interventions.

## Materials and Methods

### Animals

All animal studies were reviewed and approved by the Harvard Medical Area Standing Committee on Animal Care and Use IACUC, an AAALAC International-accredited facility. Mice were group-housed (3-4 mice per cage) at the Harvard T.H. Chan School of Public Health in standard static microisolator top cages and maintained in a temperature-controlled, pathogen-free facility with 12-h light:dark cycles with ad libitum access to autoclaved food and water. C57BL/6J mice were purchased from The Jackson Laboratory (Strain #000664). *Rosa26-LSL-TSC2-WT; Tsc2^fl/fl^* and *Rosa26-LSL-TSC2-5A; Tsc2^fl/fl^* mice were generated as previously described ^48, 88^. For liver-specific transgene expression, these lines were bred to commercially available mice expressing Albumin-Cre (Jackson Laboratory, Strain #003574)^89^. To prevent genetic drift, L-TSC2-WT and -5A colonies were maintained via heterozygous crosses of *Rosa26^LSL-TSC2-WT/LSL-TSC2-5A^; Tsc2^fl/fl^; Alb-Cre^+/-^* (L-TSC2-WT/5A) mice. To increase yield for experimental cohorts, homozygous L-TSC2-WT and -5A offspring generated from such heterozygous crosses were used as a parental generation to set up homozygous crosses. Age-matched littermates from these homozygous crosses were used for all study groups, ensuring that L-TSC2-WT and -5A lines were not bred separately for more than one generation. The *Gcgr^fl/fl^;Alb-Cre* (L-GCGR-KO) and *Gcgr^fl/fl^*mice were provided by Dr. Daniel Drucker and shared by the lab of Dr. Gökhan Hotamışlıgil^62^. L-GCGR-KO experiments utilized littermates obtained from hemizygous crosses.

To genotype mice, DNA was extracted from ear punches by the Proteinase K method, the gene regions of interest were amplified by PCR using the primer sequences below, and PCR products were analyzed by agarose gel electrophoresis. Detailed methods for genotyping TSC2-WT and - 5A mice were previously described^48^.

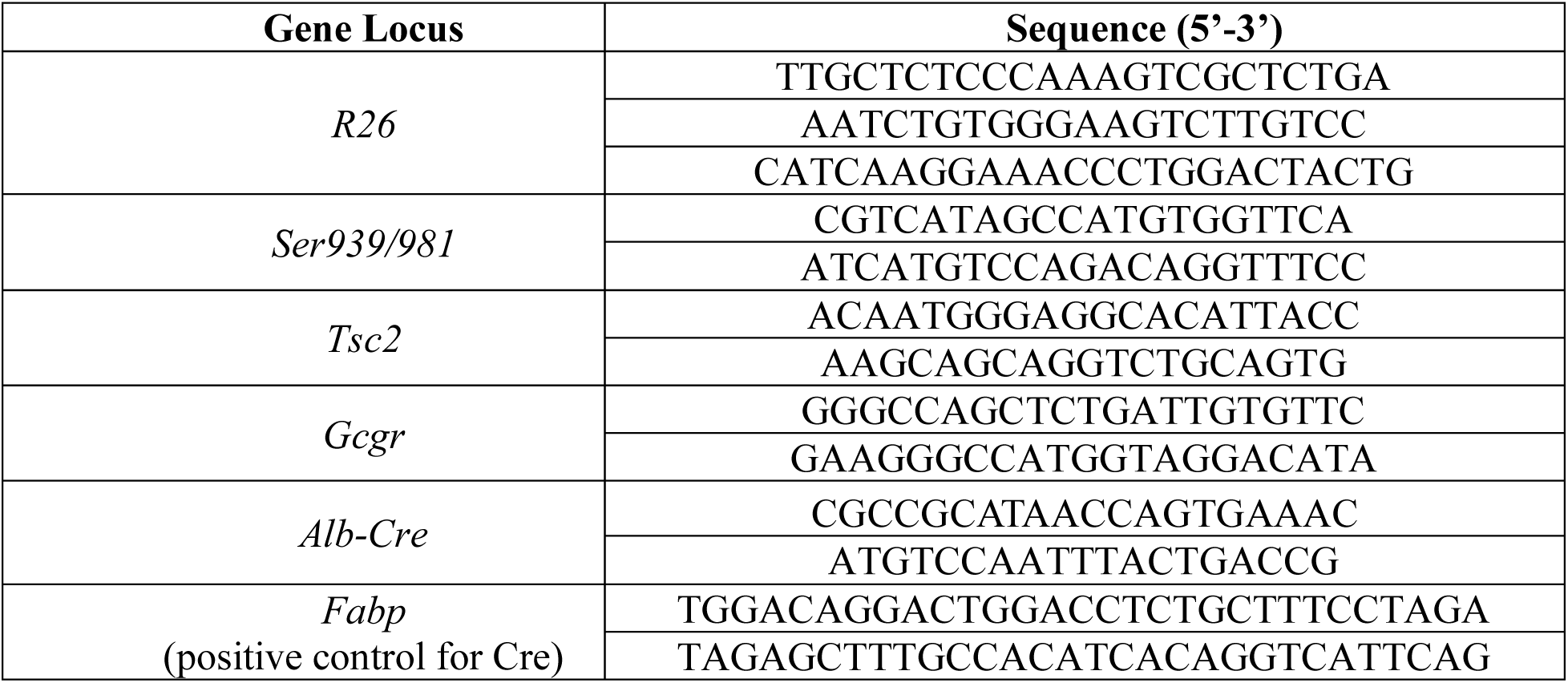

Mice were fed PicoLab Rodent Diet 20 (LabDiet, 5053) chow diet from weaning at three weeks of age. For high fat diet studies, mice were started on a 60% fat diet (Research Diets, D12492i) or a sucrose-matched control diet containing 10% fat (Research Diets, D12450Ji) beginning at 8 weeks of age. The protein-free diet was prepared using a semi-purified water-soluble diet (Research Diets, D12450BSpxi) in 1% agar (Sigma-Aldrich, #A1296), as previously described^53^. This diet mix was supplemented with casein hydrolysate (Thermo Scientific, #J12855.P5) and L-cystine (Sigma-Aldrich) for the isocaloric control diet (18% kcal protein). The high-carbohydrate diet was a variation of D12450BSpx that contained 10% kcal protein (Research Diets, D12450BSpx5i). For the protein-free or high carbohydrate diet fasting and refeeding studies, mice were acclimated to the control, semi-purified diet for 5-7 days prior to the experiment.

For animal treatments, mice were fasted overnight for 14 h followed by intraperitoneal (i.p.) injection or oral gavage of vehicle (sterile 0.9% saline), insulin (Eli Lily, HumulinR) dissolved in sterile saline with protease-free BSA at 0.5U/kg, glucose dissolved in sterile saline at 3g/kg, or feeding of the given diet, as indicated. Mice were humanely euthanized immediately prior to tissue collection, and tissues were snap-frozen in liquid nitrogen and stored at -80°C until analysis.

### Glucose, Insulin, and Pyruvate Tolerance Tests

For glucose tolerance tests, mice were fasted for 6 h during the day and injected i.p. with glucose dissolved in sterile saline at 2 g/kg (unless otherwise noted). For oral glucose tolerance tests, mice were fasted for either 6 h during the day or 16 h overnight followed by oral gavage with glucose dissolved in sterile saline at 1-2 g/kg (as indicated in figure legend). For insulin tolerance tests, mice were fasted for 6 h during the day and then injected i.p. with insulin (Eli Lily, HumulinR) in sterile saline containing 0.1% protease-free BSA at 0.5 U/kg. For the pyruvate-lactate tolerance test, mice were fasted for 16 h overnight followed by i.p. injection with lactate (1.5 mg/kg; Sigma-Aldrich, L7022) and pyruvate (150μg/kg; Sigma-Aldrich, P5280) dissolved together in sterile saline. Blood glucose was monitored using the Contour Next EZ glucometer (Bayer).

### Body composition

Body composition was measured in awake mice using the EchoMRI analyzer system (EchoMRI LLC).

### Primary Hepatocyte Isolation and Culture

Mice were anesthetized using ketamine/xylazine solution at a dose of 100 µg/10 µg per gram body weight. The liver was perfused through the portal vein with perfusion buffer (10 mM HEPES, 150 mM NaCl, 5 mM KCl, 5 mM glucose, 2.5 mM sodium bicarbonate, and 0.5 mM EDTA) for 5 minutes at a rate of 3-5 ml/min, followed by digestion buffer (10 mM HEPES, 150 mM NaCl, 5 mM KCl, 5 mM glucose, 2.5 mM sodium bicarbonate, 35 mM CaCl_2_, and 30 µg/ml Liberase TM (Sigma Aldrich)) for 5-7 minutes at a rate of 3-5 ml/min. Perfusion and digestion buffers were pre-warmed to 37°C prior to perfusion. Livers were excised and placed in DMEM (4.5g/L glucose, without glutamine or sodium pyruvate) supplemented with 2.5% FBS, 2 mM GlutaMAX, and 1% penicillin-streptomycin. Digested liver tissue was agitated to release hepatocytes, and the cell suspension was gently pelleted. Hepatocytes were purified through Percoll density separation, washed in DMEM, and seeded on collagen-coated plates (BioCoat, Corning) at a density of 1.25 x 10^6^ cells for each well of a 6-well plate. Media was replaced 4 h after seeding.

For stimulation experiments, human insulin (Sigma-Aldrich I9278) was dissolved in water and used at a concentration of 10 nM or 100 nM, glucagon (Sigma-Aldrich, G2044) was dissolved in 1% acetic acid and used at a concentration of 10 nM, forskolin (Tocris, 1099) was dissolved in DMSO and used at a concentration of 10 μM. Hepatocytes were deprived of serum overnight followed by treatment with insulin, glucagon, or forskolin for the indicated durations. For experiments in low amino acid media, standard DMEM was diluted 1:10 with filter-sterilized DMEM without amino acids (U.S. Biologicals, D-9800-27; prepared according to manufacturer’s instructions and supplemented with 4.5 g/L glucose). For amino acid stimulation, serum-starved hepatocytes were washed twice with PBS and incubated in serum-free amino-acid free DMEM for one hour, then media was replaced with standard serum-free DMEM with or without insulin.

For siRNA transfection experiments, control nontargeting pool (D-001810-10-50) and ON-Target Plus SMARTpool siRNAs against mouse *Tsc2* (L-047050-00-0020) were purchased from Horizon Discovery. Hepatocytes were transfected 4 h after seeding with 100 nM siRNA and Lipofectamine RNAiMax (6 μl/well, 6-well plate) and cultured in DMEM supplemented with 1% FBS, 1% penicillin-streptomycin, and 1 nM dexamethasone. After 72 h, the media was changed to serum-free DMEM (without dexamethasone) for insulin stimulation the following day.

### Plasma and Tissue Analysis

Plasma insulin (Crystal Chem, #90080) and albumin (Crystal Chem, #80630) were measured by ELISA following the manufacturer’s instructions. Liver glycogen was measured using the Glycogen-Glo assay (Promega, #J5051) according to manufacturer’s instructions.

### Immunoblotting and Antibodies

Primary cultured hepatocytes and liver tissue were lysed in RIPA buffer (150 mM NaCL, 5 mM EDTA, 50 mM Tis-HCl, 1% NP-40, 0.5% sodium deoxycholate, 10% SDS, 50 mM β-glycerophosphate, 50 mM NaF, 10 mM pyrophosphate tetrabasic, and 0.5mM sodium orthovanadate) with protease and phosphatase inhibitors (Sigma Aldrich). Muscle tissue was lysed in NP-40 buffer (20 mM Tris, 150 mM NaCl, 10% Glycerol, 1% NP-40) with protease and phosphatase inhibitors. Liver tissue was homogenized using the Red Lysis Kit (Next Advance) and Next Advance Bullet Blender (speed 8, for 3 min). Muscle tissue was homogenized manually using a handheld homogenizer (Cole-Parmer). Lysates were centrifuged at 20,000 x g for 15 minutes at 4°C and protein concentrations of the supernatant were determined using the Pierce Detergent-Compatible Bradford assay (Thermo Fisher Scientific). Equal amounts of protein were loaded onto polyacrylamide gels (Criterion, Bio-Rad), separated by electrophoresis, and transferred to nitrocellulose membranes. After transfer, membranes were blocked with 5% milk in Tris-buffered saline plus Tween-20 (TBST; 25 mM Tris, 150 mM NaCl, 2.5 mM KCl, 0.1% Tween-20) for 1 h at room temperature. For immunoblotting, membranes were incubated with primary antibodies (1:1000) in 5% BSA at 4°C overnight. Membranes were washed three times with TBST and then incubated with IRDye 800CW anti-Rabbit or anti-Mouse secondary antibodies (LI-COR) at 1:5000 in 5% milk for 1 h at room temperature. Membranes were washed three times in TBST and imaged using the LI-COR Odyssey CLx imaging system. LI-COR images were quantified using Image Studio Software. For Figure 7 immunoblots, membranes were developed by ECL (West Pico or Femto, Thermo Scientific) and imaged on autoradiography film. Immunoblot images on film were quantified using ImageJ software. The following primary antibodies were purchased from Cell Signaling Technologies: p-S6K1 (T389, #9234), S6K1 (#2708), p-rpS6 (S240/44; #2215), rpS6 (#2217), 4EBP1 (#9644), p-AKT (S473; #4060), AKT (#4691), p-TSC2 (T1462; #3617), TSC2 (#3612), IRS1 (#2382), p-CREB (S133, #9198), CREB (#9197), and vinculin (#4650). The β-actin antibody was purchased from Sigma-Aldrich (#A5316).

### Quantification of Plasma Amino Acids by Liquid chromatography-Mass Spectrometry

The method used for quantifying the absolute concentrations of plasma amino acids in mice following feeding was adapted from a previously described protocol^90^. Blood was collected from the tail vein in EDTA tubes (Microvette® CB 300 EDTA K2E, Sarstedt) and centrifuged to separate plasma. Plasma was mixed at 1:10 volume ratio with -20°C mixture of methanol:acetonitrile:water (40:40:20) and a stable isotope–labeled internal standard ^13^C labeled yeast extract (Cambridge Isotope Laboratory, Andover, MA, ISO1). These internal standards, together with a library of external metabolite standards, enabled absolute quantification across a wide dynamic range. The sample was vigorously vortexed and then centrifuged at 4°C at 13,000 g for 10 min. 30 μL of the resulting supernatant was transferred to an HPLC vial, from which 5 μL was injected into the LC-MS system for analysis.

Chromatographic separation was achieved using an XBridge BEH Amide XP Column (2.5 μm, 2.1 mm × 150 mm) with a guard column (2.5 μm, 2.1 mm × 5 mm) (Waters). The mobile phase consisted of water:acetonitrile 95:5 for mobile phase A and water:acetonitrile 20:80 for mobile phase B, both containing 10 mM ammonium acetate and 10 mM ammonium hydroxide. The elution linear gradient was as follows: 0–3 min, 100% B; 3.2–6.2 min, 90% B; 6.5–10.5 min, 80% B; 10.7–13.5 min, 70% B; 13.7–16 min, 45% B; and 16.5–22 min, 100% B, with a flow rate of 0.3 mL/min. The autosampler temperature was maintained at 4°C, and the injection volume was 5 μL. Needle wash was performed between samples using methanol:acetonitrile:water. The mass spectrometer used was a Q Exactive HF (Thermo Fisher), which scanned from 70 to 1000 m/z with switching polarity and a resolution of 120,000. Metabolite identification was based on accurate mass and retention time, confirmed against external standard pools. Quantification was performed either by stable isotope dilution, where a labeled internal standard was available, or by external calibration curves for metabolites lacking an internal standard. This approach enabled robust and sensitive quantification of amino acids in plasma, resolving concentrations from low nanomolar to high millimolar levels.

### Gene Expression Analysis

Liver tissue was lysed in cold TRIzol (Invitrogen) and RNA was isolated following manufacturer’s instructions. RNA was reverse transcribed using the Advanced cDNA Synthesis Kit (Bio-Rad). qPCR analysis was performed with duplicate technical replicates using iTaq SYBR green (Bio-Rad) and the CFX96 Real-Time PCR Detection System (Bio-Rad). Samples were normalized to actin (human) or 36B4 (mouse) for DDCt analysis. Primer sequences are listed below:

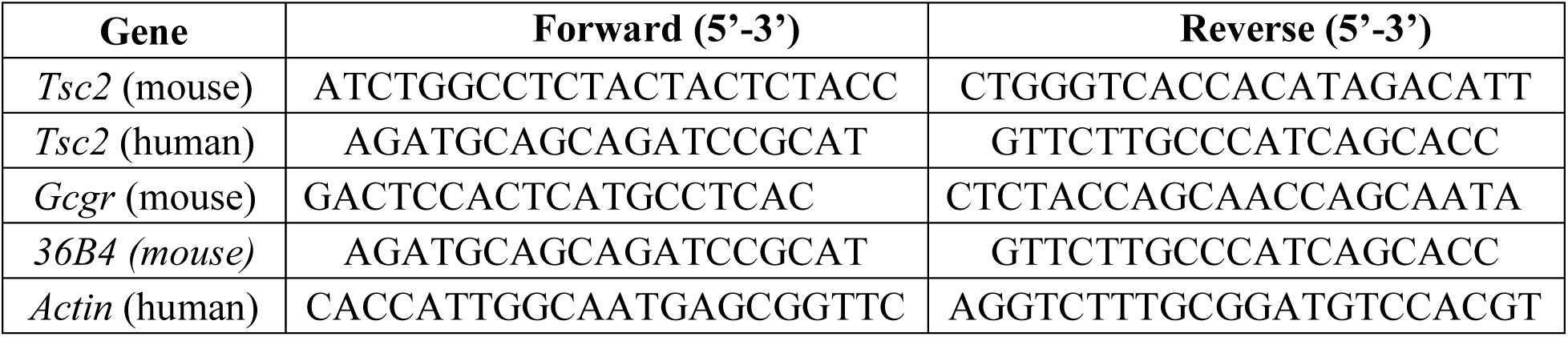

### Statistical Analysis

Statistical analyses were performed using GraphPad Prism 10.0 using appropriate tests listed at the end each figure legend. *p<0.05, **p<0.01, ***p<0.001, ****p<0.0001.

## Author Contributions

Conceptualization: K.C.K, Y.C., and B.D.M.; Investigation: K.C.K., Y.C., S.C.L, M.Y.C, and K.E.I.; Writing: K.C.K. and B.D.M.; Funding acquisition: K.C.K., Y.C., M.Y.C., and B.D.M.; Resources: G.S.H., B.D.M.; Supervision: B.D.M.

## Acknowledgements

We thank Vanessa Byles for her insights on liver biology and primary hepatocyte culturing; Julissa Reyes for technical assistance with animal care; Bo Yuan, Juliya Hsiang, and Sheng Tony Hui of the The Harvard Chan Advanced Multi-omics Platform (ChAMP) for mass spectrometry sample analysis; and Michael MacArthur and Sarah Mitchell for advice on dietary formulation. We thank Chih-Hao Lee, Sudha Biddinger, Pere Puigserver, and members of the Manning lab for their scientific insight and critical feedback. Additional thanks to Daniel Drucker for kindly sharing the *Gcgr^fl/fl^* and L-GCGR-KO mice. This study was supported by a Glenn Foundation for Medical Research Postdoctoral Fellowship (Y.C.); NIH grants T32-DK128781 (K.C.K. and S.C.L.), F31-DK128873 (K.C.K.), R35-CA197459 (B.D.M.), and P01-CA120964 (B.D.M.); and Damon Runyon Cancer Research Foundation Merck Fellowship DRG-#2443-21 (M.Y.C.). These funders were not involved in the design, execution, or interpretation of this study.

## Notes

### Competing Interest Statement

B.D.M. is a consultant for Gondoka Bio independent of the contents of this article. G.S.H. is a member of the Scientific Advisory Board and holds equity in Crescenta Pharmaceuticals (not related to the contents of this article). Additionally, technology from the Hotamışlıgil Lab is licensed to Iş Private Equity, from which G.S.H. also receives research funding. This is also not related to the contents of the article. All the other authors declare no competing interests.

