## Supplemental Figures for "Dietary protein governs the role of insulin signaling in the postprandial regulation of hepatic mTORC1"

#### Supplemental Figure Legends

**Figure S1. Phenotypic characterization of L-TSC2-WT and -5A females.**

(A) Liver weights of female L-TSC2-WT and -5A mice fed ad libitum at 6 months of age graphed as mean ± SD; n = 8 WT, 13 5A.

(B) Body weights of female L-TSC2-WT and -5A mice up to 6 months of age graphed over time as mean ± SD; n=6-13/group/timepoint. There were no significant differences in body weight between L-TSC2-WT and -5A mice at any time point tested.

(C, D) Fat (C) and lean (D) mass of L-TSC2-WT and -5A female mice measured by EchoMRI at indicated ages and graphed as mean ± SD; n = 6-13/group.

(E) Plasma insulin following 6-h daytime fast in 6-month-old female L-TSC2-WT and -5A mice graphed as mean ± SD; n = 8 WT, 105A.

(F) Glucose tolerance test performed on 6-month-old L-TSC2-WT and -5A female mice after a 6-h daytime fast followed by i.p. injection of glucose (2 g/kg). Blood glucose measurements over time are plotted as mean ± SD. AUC values are graphed as mean ± SEM; n = 8 WT, 10 5A.

(G) Insulin tolerance test performed on 6-month-old L-TSC2-WT and -5A female mice after a 6-h daytime fast followed by i.p. injection of insulin (0.5 U/kg). Blood glucose measurements over time are plotted as mean ± SD. AUC values are graphed as mean ± SEM; n=8 WT, 10 5A.

Statistical analysis: (A and E) Welch's t test; (B) mixed-effects analysis with Šidák correction; (C and D) two-way ANOVA with Šidák correction; (F and G) two-way ANOVA (F) or mixed-effects analysis (G) with Šidák correction for curves, Welch's t-test for AUC. ns $p \geq 0.05$.

**Figure S2. mTORC1 activation is disconnected from AKT-TSC2 regulation specifically in the liver of L-TSC2-5A mice with no changes to insulin induction.**

(A) Immunoblots of gastrocnemius lysates from 11-week-old L-TSC2-WT or -5A male mice fasted for 14 h overnight followed by i.p. injection with vehicle (saline) or insulin (0.5 U/kg) for 20 min with accompanying bar graphs of the mean ± SD of phospho-to-total protein signal quantifications normalized to the L-TSC2-WT vehicle-treated group; n = 5/group.

(B) Blood glucose measured in fasted L-TSC2-WT and -5A male mice 25 min after i.p. injection with vehicle (saline) or glucose (3 g/kg) graphed as mean ± SD; n = 4 WT, 5 5A.

(C) Plasma insulin measured in blood collected from fasted L-TSC2-WT and -5A male mice 25 min after i.p. injection with vehicle (saline) or glucose (3 g/kg) graphed as mean ± SD; n = 4 WT, 5 5A.

(D) Plasma insulin measured in blood collected from L-TSC2-WT and -5A male mice fasted overnight for 14 h or fasted and then refed for 1 h graphed as mean ± SD; n = 3-4/group.

Statistical analysis: (A-D) two-way ANOVA with Šidák correction. ns $p \geq 0.05$, *$p < 0.05$, **$p < 0.01$, ***$p < 0.001$, and ****$p < 0.0001$.

**Figure S3. Dietary protein (or amino acids) controls insulin responsiveness of hepatic mTORC1 activation.**

(A) Blood glucose values in L-TSC2-WT and -5A male mice fasted for 14 h overnight and, where indicated, 30 min after refeeding a semi-purified control (CTL) diet or a protein-free (PF) diet graphed as mean ± SD; n = 10/group fasted, 5/group refed.

(B) Phospho-to-total S6K signal from primary mouse hepatocytes treated with an insulin (10 nM) time course under high amino acid conditions graphed as mean ± SD fold induction from the L-TSC2-WT vehicle condition (calculated separately for each replicate). The notations directly above each bar indicate whether the signal was significantly induced from its respective vehicle control for each genotype; n = 3 biological replicates.

(C-F) Male L-TSC2-WT or -5A mice were fed a high carbohydrate, low protein (HC-LP) diet starting at 8 weeks of age.

(C) Body weight of L-TSC2-WT and -5A male mice after 6 weeks HC-LP diet feeding graphed as mean ± SD; n = 11 WT, 12 5A.

(D, E) Fat (D) and lean (E) mass of L-TSC2-WT and -5A male mice after 6 weeks HC-LP diet feeding measured by EchoMRI and graphed as mean ± SD; n = 11 WT, 12 5A.

(F) Plasma insulin of L-TSC2-WT and -5A male mice fed the HC-LP diet for 5 weeks and fasted for 16 h overnight graphed as mean ± SD; n = 11 WT, 12 5A.

Statistical analysis:(A and B) two-way ANOVA with Šidák correction; (C-F) Welch's t-test. ns $p \geq 0.05$, *$p < 0.05$, **$p < 0.01$, ***$p < 0.001$, and ****$p < 0.0001$.

**Figure S4. Muscle mTORC1 activation is elevated in obese mice.**

(A-E) Male C57BL/6J mice were fed a high-fat diet (HFD) or low-fat control diet (LFD) beginning at eight weeks of age.

(A, B) Fat (A) and lean (B) mass of C57BL/6J male mice at eight weeks of age and after 4 or 8 weeks of HFD or LFD feeding measured by EchoMRI and graphed as mean ± SD; n = 18 for baseline, n=6 for 4 or 8 weeks.

(C, D) Glucose tolerance test performed after 4 (C) and 8 (D) weeks HFD or LFD. Mice were fasted for 6 h followed by i.p. injection of glucose (2 g/kg). Blood glucose measurements over time are plotted as mean ± SD. AUC values are graphed as mean ± SEM; n = 6/ group.

(E) Immunoblots of gastrocnemius lysates from mice fed HFD or LFD for 8 or 15 weeks and fasted for 6 h (daytime) prior to tissue collection. Accompanying bar graphs of phospho-to-total protein signal quantifications normalized to the LFD group plotted as the mean ± SD are shown; n = 6/group.

Statistical analysis: (A and B) two-way ANOVA with Šidák correction; (C-D) two-way ANOVA (C) or mixed-effects analysis (D) with Šidák correction for curves, Welch's t-test for AUC; (E) Mann-Whitney test. ns $p \geq 0.05$, *$p < 0.05$, **$p < 0.01$, ***$p < 0.001$, and ****$p < 0.0001$.

**Figure S5. L-TSC2-5A mice are not protected from metabolic dysregulation in response to diet-induced obesity.**

(A-C) Male L-TSC2-WT and -5A mice were fed high-fat diet (HFD) beginning at eight weeks of age.

(A) Plasma insulin following 6 h fasting in L-TSC2-WT and -5A male mice fed HFD for 0, 4, 8, and 16 weeks graphed as mean ± SD; n = 8-9 WT, 8-12 5A.

(B) Glycogen content in liver tissue collected from ad libitum-fed L-TSC2-WT and -5A male mice fed HFD for 20 weeks graphed as mean ± SD; n = 7 WT, 9 5A.

(C) Albumin concentration in plasma collected from ad libitum-fed L-TS2-WT and -5A male mice fed HFD for 20 weeks graphed as mean ± SD; n = 7 WT, 10 5A.

(D-I) Female L-TSC2-WT and -5A mice were fed high-fat diet (HFD) beginning at eight weeks of age.

(D) Body weight measurements of L-TSC2-WT and -5A female mice fed HFD for indicated amount of time graphed as mean ± SD; n = 8 WT, 10-13 5A.

(E, F) Fat (E) and lean (F) mass of L-TSC2-WT and -5A female mice at baseline (eight weeks of age) and after 8 or 16 weeks HFD measured by EchoMRI and graphed as mean ± SD; n = 6 WT, 9 5A.

(G) Plasma insulin measurements in blood collected from L-TSC2-WT and -5A females fasted for 6 h following 16 weeks HFD graphed as mean ± SD; n = 7 WT, 9 5A.

(H) Glucose tolerance test performed in L-TSC2-WT and -5A females after 15 weeks HFD. Mice were fasted for 6 h followed by i.p. injection of glucose (2 g/kg). Blood glucose measurements over time are plotted as mean ± SD. AUC values are graphed as mean ± SEM; n= 6 WT, 9 5A.

(I) Insulin tolerance test performed in L-TSC2-WT and -5A females after 16 weeks HFD. Mice were fasted for 6 h followed by i.p. injection with insulin (0.5 U/kg). Blood glucose measurements over time are plotted as mean ± SD. AUC values are graphed as mean ± SEM; n = 6 WT, 9 5A.

Statistical analysis: (A, E and F) two-way ANOVA with Šidák correction; (B and C) Welch's t-test; (D) mixed-effects analysis with Šidák's correction; (G) Welch's t-test; (H and I) two-way ANOVA (H) or mixed-effects analysis (I) with Šidák correction for curves, Welch's t-test for AUC. ns $p \geq 0.05$.

### Figure S1

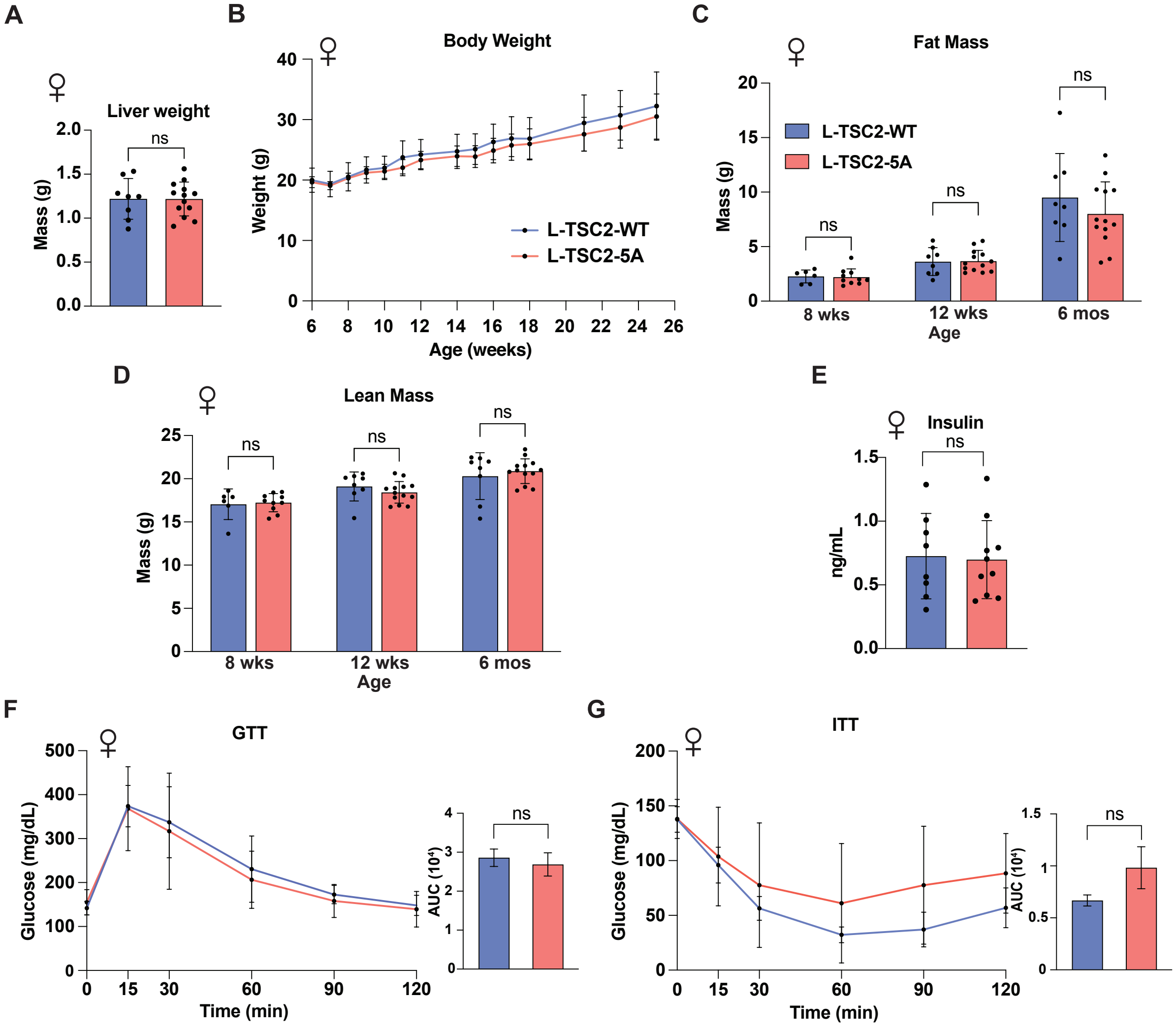
A
Liver weight
Mass (g)
ns
B
Body Weight
Weight (g)
Age (weeks)
L-TSC2-WT
L-TSC2-5A
C
Fat Mass
Mass (g)
L-TSC2-WT
L-TSC2-5A
ns
8 wks
12 wks
6 mos
Age
D
Lean Mass
Mass (g)
ns
8 wks
12 wks
6 mos
Age
E
Insulin
ng/mL
ns
F
GTT
Glucose (mg/dL)
Time (min)
AUC (10⁴)
ns
G
ITT
Glucose (mg/dL)
Time (min)
AUC (10⁴)
ns

### Figure S2

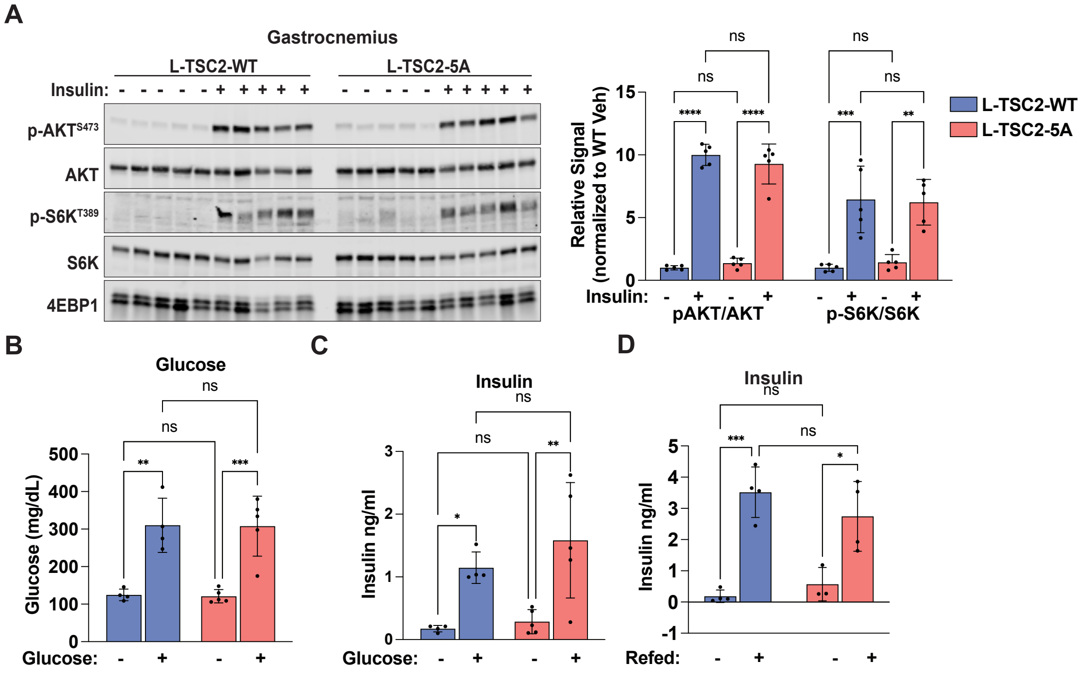
A
Gastrocnemius
L-TSC2-WT
L-TSC2-5A
Insulin:
p-AKT S473
AKT
p-S6K T389
S6K
4EBP1
Relative Signal (normalized to WT Veh)
pAKT/AKT
p-S6K/S6K
L-TSC2-WT
L-TSC2-5A
B
Glucose
Glucose (mg/dL)
Glucose:
C
Insulin
Insulin ng/ml
Glucose:
D
Insulin
Insulin ng/ml
Refed:

Figure S3

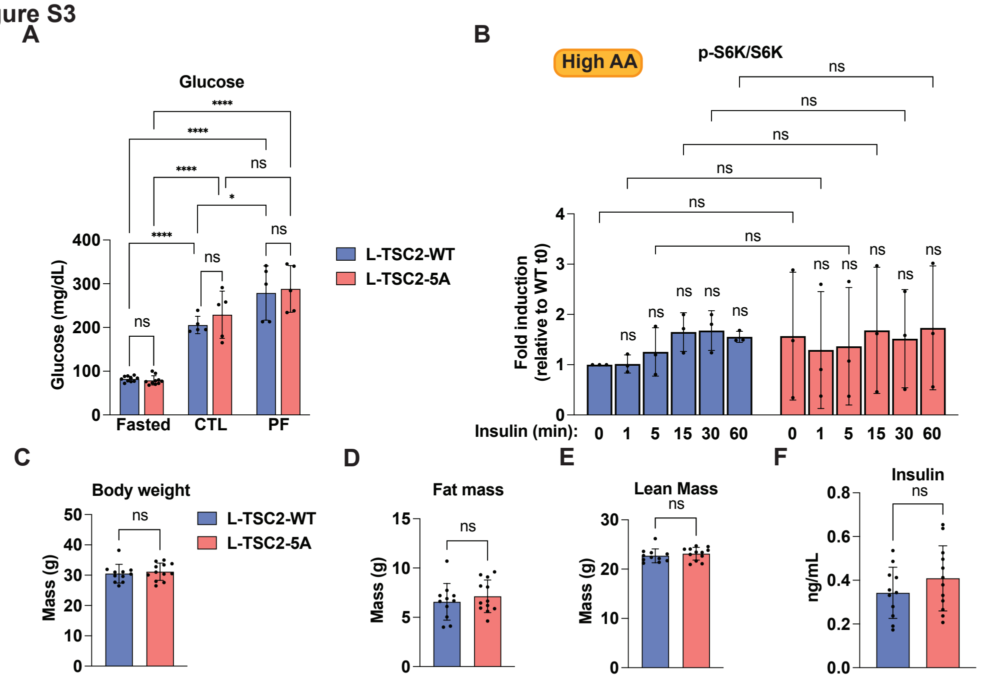
A
Glossary
Glucose
Glucose (mg/dL)
Fasted
CTL
PF
L-TSC2-WT
L-TSC2-5A
ns
****
*
B
High AA
p-S6K/S6K
Fold induction (relative to WT t0)
Insulin (min):
0
1
5
15
30
60
C
Body weight
Mass (g)
D
Fat mass
E
Lean Mass
F
Insulin
ng/mL

Figure S4

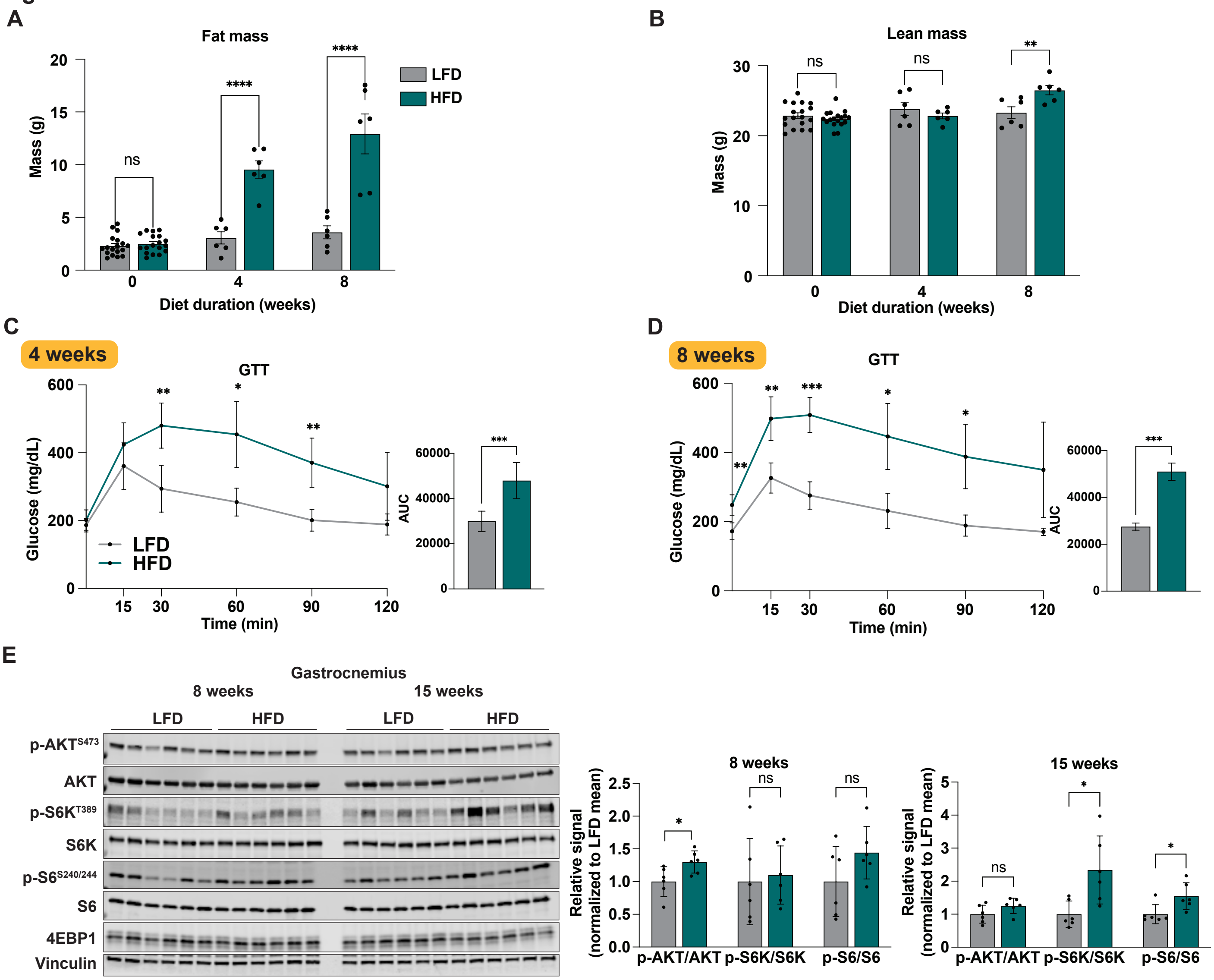
A
Fat mass
Mass (g)
Diet duration (weeks)
LFD
HFD
B
Lean mass
C
4 weeks
GTT
Glucose (mg/dL)
Time (min)
AUC
D
8 weeks
E
Gastrocnemius
8 weeks
15 weeks
p-AKTS473
AKT
p-S6KT389
S6K
p-S6S240/244
S6
4EBP1
Vinculin
Relative signal (normalized to LFD mean)
p-AKT/AKT
p-S6K/S6K
p-S6/S6

Figure S5

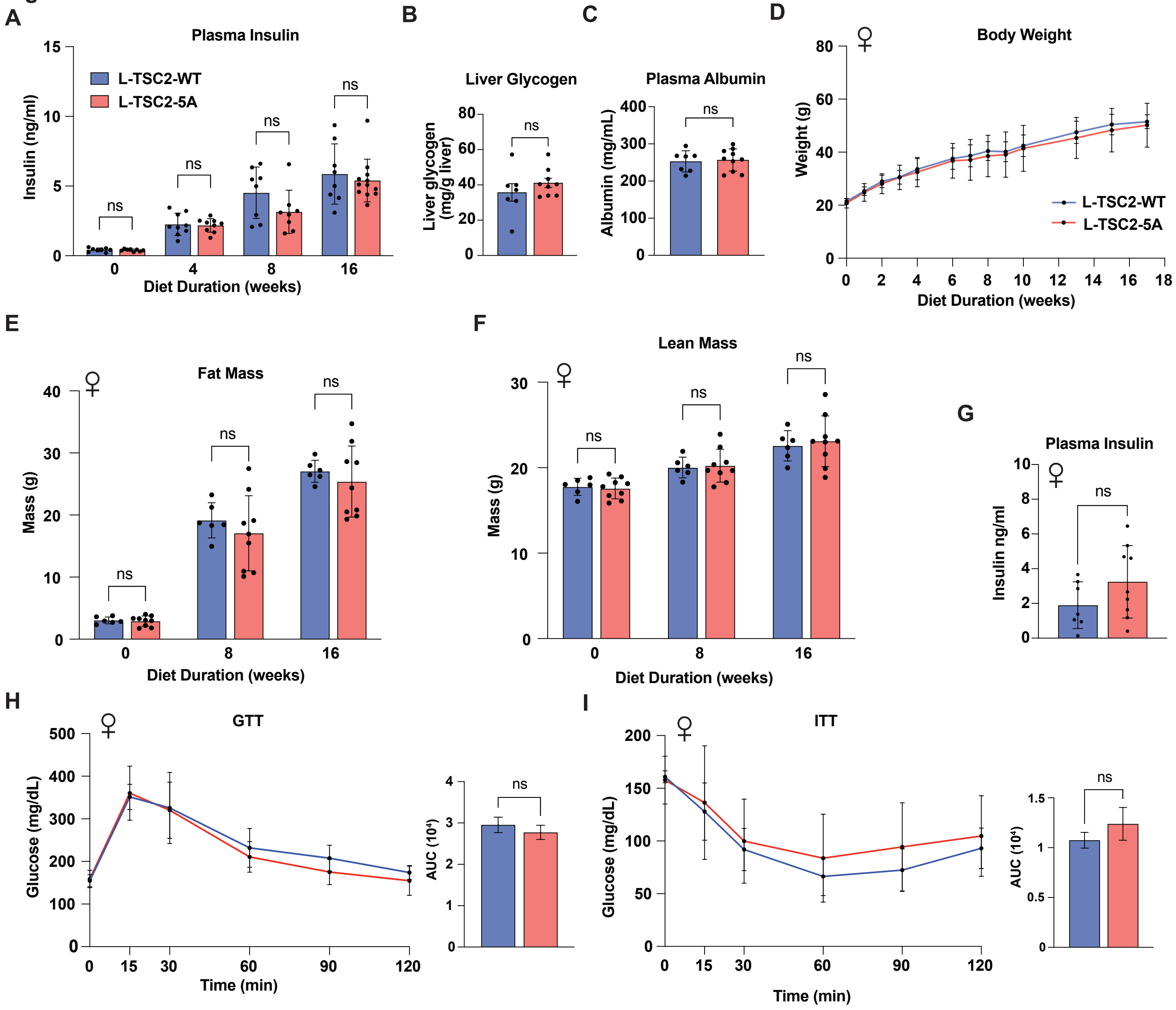
A
Plasma Insulin
L-TSC2-WT
L-TSC2-5A
Insulin (ng/ml)
Diet Duration (weeks)
ns
B
Liver Glycogen
Liver glycogen (mg/g liver)
C
Plasma Albumin
Albumin (mg/mL)
D
Body Weight
Weight (g)
Diet Duration (weeks)
E
Fat Mass
Mass (g)
F
Lean Mass
G
Plasma Insulin
Insulin ng/ml
H
GTT
Glucose (mg/dL)
Time (min)
AUC (10⁴)
I
ITT
